# A novel cell indirect calorimetry method unveils the metabolic fluxomic signatures of human monocyte derived M(LPS+INF-γ) and M(IL-4) macrophages

**DOI:** 10.1101/2025.10.21.681401

**Authors:** Gloria Cinquegrani, Valentina Spigoni, Federica Fantuzzi, Anna D’Antuono, Francesca Bagnaresi, Elisabetta Giordano, Alberto Burato, Alessandra Dei Cas, Riccardo C. Bonadonna

## Abstract

Macrophages (MΦ) display distinct immunometabolic phenotypes upon polarization. While transcriptomic analyses have suggested divergent metabolic programs in human M(LPS+INF-γ) and M(IL-4) MΦ, a comprehensive assessment of their metabolic fluxes is lacking.

**Aim** of this study is to 1. develop and validate a novel indirect microcalorimetry method for quantifying cellular metabolic fluxes, and 2. exploit it to characterize fluxomic signatures of polarized human monocyte-derived macrophages.

**Methods:** MΦ from healthy donors were differentiated into M0, M(LPS+INF-γ), and M(IL-4) phenotypes and studied in four defined media: substrate-free, glucose, glycyl-glutamine, and glucose + glycyl-glutamine. A steady-state fluxomic model was constructed by integrating four independent measures – oxygen consumption and proton production (Seahorse XFp), lactate and ammonia release (microfluorimetry) – into stoichiometric equations of metabolism (SAAM II software).

**Results:** Fluxes revealed that macrophages rely on glucose to sustain glycolysis, contributing ∼30% of citrate synthase flux, and predominantly on lipids for net citrate synthesis (first step of Krebs cycle). Upon polarization, M(LPS+INF-γ) macrophages showed increased anaerobic glycolysis versus M0 and M(IL-4), with similar TCA fluxes to M0. In contrast, M(IL-4) macrophages exhibited higher TCA and malic enzyme fluxes, especially with glucose and glycyl-glutamine, and a trend toward enhanced lipid oxidation.

**Conclusions:** This novel method enables precise quantification of bioenergetic fluxes. In human MΦ, it reveals that M(LPS+INF-γ) and M(IL-4) subsets exhibit distinct metabolic phenotypes, consistent with their immunological roles. These results resolve transcriptomic-metabolic discrepancies and provide a robust framework for assessing immunometabolism in primary human cells.

## Introduction

Metabolism is a core process underlying essentially all biological processes, as cells require energy to sustain growth, differentiation and physiological activities. The investigation of the crosstalk between metabolic changes and immune cell function has unveiled metabolic fingerprints underlying immune system (dys)function in health and disease (1). Thus, cell metabolism assessment is essential for understanding and defining (immune) cell functions and in helping to diagnose certain human diseases (2). This general concept has been encapsulated in the term “immunometabolism”.

A paradigmatic example of immunometabolism is embodied by macrophages (MΦ). These cells are very plastic and modify their gene expression and transcription profile along a continuous spectrum, with previously named M1 and M2 MΦ phenotypes as extremes, in response to changes in their environment (3). The microbial component lipopolysaccharide (LPS) can drive macrophage polarization to the pro-inflammatory M1 phenotype, while interleukin 4 (IL-4) can induce macrophage polarization to immunosuppressive M2a. In our previous work, we demonstrated by transcriptomic analysis *in vitro* that human M(LPS+INF-γ) *vs* M(IL-4) are featured by distinct bioenergetic pathways. Specifically, GSEA analysis of M(LPS+INF-γ) activation pointed out a fall in the expression of the genes of mitochondrial metabolism and cellular respiration, whereas M(IL-4) activation showed the opposite pattern (4). This was reminiscent of the immunometabolic evidence obtained in murine MΦ, in which M(LPS+INF-γ) primarily rely on glucose/glycogen utilization and anaerobic glycolysis and M(IL-4) exploit oxidative phosphorylation (OXPHOS) and fatty acid oxidation for ATP supply (5, 6). Thus, M(LPS+INF-γ) and M(IL-4) phenotypes are thought to feature divergent metabolic substrate preference and pathway activation, turning out as an excellent model for immunometabolic studies (7). The immunometabolic pattern of murine MΦ may not be directly superimposable to human MΦ. According to most recent evidence, human monocyte derived pro-inflammatory MΦ display accelerated glycolysis and a concomitant reduction in oxidative metabolism, whereas M(IL-4) decrease glycolysis with little or no change in oxidative metabolism (8, 9).

The discrepancy between transcriptomic data and metabolic fluxes, especially in M(IL-4), may be underlied primarily by one or both of two different factors: i. transcriptomics, albeit fundamental, is just gene transcription and cannot substitute for the assessment of real metabolic fluxes, a limitation shared also by other techniques – e.g. single-cell RNA sequencing (scRNA-seq) (10) and metabolomics; ii. the assessment of metabolic fluxes in the human studies focused on MΦ metabolism may have been fraught with limitations.

Following our experience with transcriptomics of human M(LPS+INF-γ) and M(IL-4) (4), we turned to build an experimental setting to apply fluxomics (11), which first of all requires to measure the rates (fluxes; mass/min) of exchange/transfer of molecules in a biological reactor, the latter varying in complexity from subcellular systems up to entire, intact organisms. As thoroughly discussed in (10), it is rather common to find discrepancies between transcriptomic/proteomic data and measured fluxes in central carbon metabolism. Thus, primary measures of the metabolic fluxes, albeit not exhaustive and wanting integration with “omics” data, should be central to quantify cell metabolism (10).

As to bioenergetic metabolism and energy turnover, which are thought to be the metabolic signature of MΦ subtypes, the assessment of both glycolysis and mitochondrial respiration is of paramount importance (2). For these purposes, there is widespread use of technologies, which provide a continuous recording of oxygen consumption rate (OCR) and extracellular acidification rate (ECAR) of viable cells, without the use of dyes or tracers (12). This approach is widely used to gain insights in some core metabolic pathways, but it relies on a series of assumptions, the relevance and the limitations of which not always are properly appreciated. For instance, in most cases it is given for granted that ECAR or H^+^ production rate is a reliable *proxy* of cell lactate production, thereby assuming that proton generation by a plethora of metabolic reactions, including, but not limited to, CO_2_ production by the tricarboxylic acid (TCA) cycle, would not deserve to be formally taken into account (13, 14).

Mookerje *et al.* exploited the measures of OCR and ECAR, the latter one after conversion to a *bona fide* proton production rate (PPR), to improve the estimates of both respiration and glycolysis contribution to extracellular H^+^ production (15) and to quantify intracellular ATP turnover (16).

However, these models included only glycogen and/or exogenous glucose as cell energetic sources, neglecting, or a priori assuming, lipid and/or amino acid contributions to energetic metabolism.

Thus, although Mokerjee et al.’s work was, and is, a huge step forward in the assessment of cell bioenergetic fluxes, perhaps there might still be significant room for improvement.

In this paper, our first goal was to extend Mokerjee’s et al. work by introducing a novel technique of indirect microcalorimetry to quantify cell bionergetic fluxes. In analogy to indirect calorimetry in the intact organism (17, 18), the cell indirect microcalorimetry technique herein introduced exploits few primary measurements (namely OCR, PPR, lactate and ammonia release rates) to calculate the main cell bioenergetic fluxes under steady state conditions, by a system of equations based on the stoichiometry of known biochemical reactions. A preliminary validation was searched by investigating the metabolic response of MΦ metabolism to different substrates.

As a second goal of this paper, we applied this newly developed cell indirect calorimetry method to clarify whether or not human monocyte derived M(LPS+INF-γ) MΦ and M(IL-4) MΦ truly display catabolic/energy metabolism rates which are in contradiction with their previously reported transcriptomic signatures (10).

## Methods

### Ethics Statement

The local Institutional Review Board - Comitato Etico di Area Vasta Emilia Nord - approved the study protocol (prot. n. 22597), which was conducted in accordance with the Declaration of Helsinki. Peripheral blood mononuclear cells (PBMCs) were obtained from healthy donors who gave written informed consent.

### Macrophage culture and experimental conditions

Macrophages (MΦ) were isolated from healthy donors’ buffy-coats and cultured as previously reported (4, 19). Briefly, PBMCs were isolated by Lymphoprep density gradient centrifugation (Euroclone) and seeded in six-well plates at a density of 2×10^7^ cells/ml. Monocytes were selected by plastic adherence for 1h and cultured for 6 days at 37°C and 5% CO_2_ in RPMI 1640 medium (Euroclone) supplemented with 10% FBS, 1% L-glutamine, 1% pen/strep, 1% amphotericin B and 70 ng/ml Macrophage-Colony Stimulating Factor (M-CSF) (Miltenyi Biotech, Bergisch Gladbach, Germany). Polarization was obtained by stimulating MΦ for 16 h with LPS (100 ng/ml) and IFN-γ (20 ng/ml) (both purchased by Miltenyi Biotech) for M(LPS+IFN-γ) MΦ (previously named M1); and with IL-4 (20 ng/ml; Miltenyi Biotech) for M(IL-4) MΦ (previously named M2a). No stimuli were added in culture to obtain resting MΦ (M0), as a control condition.

### Inflammatory profile of human polarized macrophages

To assess MΦ inflammatory profile and phenotype, we performed quantitative PCR (qPCR) assays of candidate genes. Briefly, cells were lysed by Qiazol and total RNA was isolated by miRNeasy mini kit (Qiagen Ltd, West Sussex, UK) followed by NanoDrop (Thermo Fisher Scientific, Waltham, Massachusetts, USA) quantification and quality check. A total of 300 ng of RNA were reverse transcribed using High-Capacity RNA-to-cDNA Kit (Applied Biosystem, Life Technologies, Foster City, California, USA), following manufacturer’s instructions.

The gene expression of CD80 (Hs.PT.56a.38577902), CD200R1 (Hs.PT.58.3206692), CD206 (Hs.PT.58.15093573), CD209 (Hs.PT.58.40094997) all purchased from Integrated DNA Technologies (Coralville, Iowa, USA), and of interleukin (IL)-6 (Hs00174131_m1), IL-8 (Hs00174103_m1), IL-1β (Hs01555410_m1), Tumor Necrosis Factor (TNF)-α (Hs00174128_m1; all from Applied Biosystems) was assessed using TaqMan Universal Master Mix (Applied Biosystems) with TaqMan primers and probes on a CFX Connect Real-Time (Bio-Rad, Hercules, CA, USA), as previously reported (20, 21). Thermal cycling conditions were as follows: 98° for 30s, followed by 40 amplification cycles (95°C for 15s; 60°C for 1 min).

Gene expression values were normalized to the geometric mean of ribosomal protein S18 (RPS18) (Hs01375212_g1), glyceraldehyde 3-phosphodehydrogenase (GAPDH) (Hs03929097_g1), β-actin (ACTB) (Hs99999903_m1), β-2-Microglobulin (B2M) (Hs00187842_m1) reference genes. Each sample was analysed in triplicate and the mean values were used for calculations based on the ΔΔCt method (22).

### Macrophage phenotypic characterization

To confirm the phenotypes of polarized MΦ, 1×10⁶ cells were analysed by flow cytometry (LSRFortessa, BD Biosciences, Franklin Lakes, NJ, USA) for the surface expression of CD80 (BB515 Mouse Anti-Human CD80) and CD209 (PE Mouse Anti-Human CD209) (both from BD Biosciences). To exclude non-viable cells, samples were stained with 7-AAD (7-aminoactinomycin D; BD Biosciences). Cells negative for 7-AAD staining (i.e., viable cells) were gated and analysed for CD80 and CD209 expression, reported as mean fluorescence intensity (MFI) values. MFI values were corrected for autofluorescence by subtracting the MFI of the unstained control from that of the stained sample. For each analysis, at least 1×10⁵ events were acquired using FACSDiva software.

All flow cytometric assays and data analyses were performed by a single trained operator.

### Macrophage handling for metabolic assessment

On day 6, unstimulated MΦ were harvested and re-plated in 80 µl of RPMI at 5×10^4^ cells/well in uncoated Seahorse Flux microplates. MΦ were incubated with polarization stimuli for 16 h in standard culture conditions. Each experimental condition was performed in duplicate and fluxomic analysis was performed on each well, separately. Then, the mean value of each fluxomic parameter relating to the specific culture condition was considered for bioenergetic calculations.

### Seahorse Assay procedures

One hour before the assay run, cells were washed three times and then incubated in 180 µl of KREBS medium (2 mM HEPES, 136 mM NaCl, 2 mM NaH_2_PO_4_, 3.7 mM KCl, 1 mM MgCl_2_, 1.5 mM CaCl_2_, 500 U/ml carbonic anhydrase, 0.1% (w/v) fatty-acid-free BSA), at 37 °C in a non-CO_2_ incubator, under four different substrate supply conditions: i. no substrates; ii 5 mM glucose; iii. 2 mM glycyl-glutamine (GlyGln); iv. 5 mM glucose + 2 mM GlyGln.

KREBS buffer was preferred to DMEM because it lacks any metabolic substrate (i.e. amino acids), red phenol and bicarbonate buffer. We also noticed that culture media enriched with amino acids or glutamine alone spontaneously release significant amounts of NH_3_ + H^+^, thereby banning the use of NH_4_ as a readout of irreversible protein/amino acid catabolism by the cultured cells (see below). We used glycyl-glutamine (GlyGln) in place of glutamine, because it is stable in solution and it can be readily utilized by cells as an oxidative substrate, contributing to the pyruvate pool via glycine and to the oxaloacetate pool through the entry of the glutamine carbon skeleton as α-ketoglutarate into the TCA cycle. Importantly, since it does not undergo significant spontaneous deamination, GlyGln enables the use of NH_4_ as a readout of protein/amino acid catabolism by the cells.

### Seahorse-based assay

To perform cellular indirect calorimetry under different culture conditions, baseline values of oxygen consumption rate (OCR) and extracellular acidification rate (ECAR) were recorded using the Seahorse XF Analyzer. These baseline measurements were obtained from three consecutive measurement cycles, each consisting of a 1-minute mixing phase, a 1-minute wait phase, and a 3-minute measurement phase. Please, note that the time of the run (into the seahorse instrument) is of about 35 min, but cells were incubated in KREBS medium for overall 110 min. No stimulus/modulator was injected during the assays. For calculation, the last OCR and ECAR recorded value was used, following parameter stabilization. All OCR and ECAR data were normalized to the number of viable cells.

To allow conversion of the ECAR value (mpH/min) to proton production rate (PPR; pmol H^+^/min), which is the metabolic flux of interest, we measured the buffering factor of KREBS medium by performing a titration with HCl (Seahorse XFp Analyzer, Agilent Technologies) for each medium used (substrate deprived medium, glucose, glycyl-glutamine and glucose plus glycyl-glutamine), as described in (14, 21). Briefly, during the assay the Seahorse instrument injected four sequential HCl (0.2 mM) volumes, leading to a progressive decrease in pH. The pH variations were recorded, and differences (ΔpH) were calculated after each injection. Since pH responded pseudo-linearly to H^+^, a regression line between ΔpH and H^+^ nmol of the measuring volume (2.28 µL) was calculated. The slope of the regression line was the buffering factor (BF) of the medium.

#### Mitostress test

In four donors, mitochondrial function of M0, M(LPS+IFN-γ) and M(IL-4) in glucose plus glycyl-glutamine medium was tested by Cell Mito Stress Test (Seahorse XFp Analyzer; Agilent Technologies), following the protocol provided by the manufacturer. The assay measured OCR in cells after sequential injections of 1.5 μM oligomycin, 1 μM FCCP, and 1 μM rotenone+antimycin A to calculate basal and ATP-linked respiration, proton leak, maximal respiration, spare respiratory capacity, and non-mitochondrial respiration. All OCR values were normalized to a fixed cell number (5 x 10^4^), as determined by cell normalization assay. These experiments were conducted to test whether different proton leak rates were present in the three MΦ subtypes.

### Cell normalization assay

To normalize OCR and PPR data, cell number was determined by CyQuant Cell Proliferation Assay kit (Thermo Fisher Scientific) which measures double-stranded DNA in each microwell. Briefly, immediately after the Seahorse assay, supernatants were collected and microplates stored in ultra-freezer. The day of the assay, 200 µl of CyQuant GR working solution (1:20 cell-lysis buffer + 1:400 CyQuant GR stock solution) were added to each well. After incubation (5 min at room temperature), fluorescence was measured (excitation: 480 nm, emission: 520 nm) by Varioskan LUX (Thermo Fisher Scientific). Cell number was calculated by using a standard curve generated by serially 2-fold diluting a fixed amount of cells.

### Supernatant deproteinization

Following Seahorse assays, supernatants were collected and deproteinized to remove potential traces of lactate dehydrogenase. After thawing, samples were diluted 1:10 in metaphosphoric acid (MPA) 2 mol/l and centrifuged at 10.000 g for 5 min at 4°C. Then 1:12 of KHCO_3_ 2 mol/l was added to the collected supernatants and centrifuged again at 10.000 g for 5 min at 4°C. Deproteinized supernatants were stored at-80°C for NH_4_ and lactate quantification.

### Metabolite assays

The Lactate Colorimetric/Fluorometric Assay Kit (BioVision) was used to quantify lactate release by MΦ in deproteinized media, following manufacturer’s instructions with some changes. The final volume of the reaction was adapted to 40 μl as follows: 0.5 μl of samples (diluted 1:40 in assay buffer to a volume of 20 μl) and 20 μl of reaction mix (19 μl of assay buffer plus 0.8 μl of enzyme mix and 0.2 μl of probe). After 15 min of incubation at room temperature, the fluorescent signal (excitation: 535 nm, emission: 595 nm) was measured by a Varioskan LUX (Thermo Fisher Scientific). Sample concentration was measured by using a standard curve [40 - 200 pmol/well; Limit of detection (LoD)=20 pmol/well] and expressed as pmol/min in relation to assay duration.

To minimize differences with samples, all standard curve points were prepared in KREBS medium and deproteinized. Each sample was analysed in duplicate and the mean value was used in fluxomic equations.

Ammonia (NH_4_) concentration in deproteinized supernatants was assessed by Ammonia Assay kit (Merck, Darmstadt, Germany). The assay is based on this specific reaction (eq. 1):

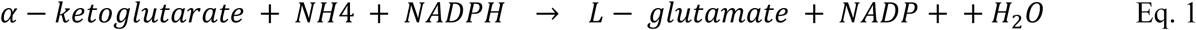

This fluorometric assay is based on the reduction of RFU values due to the oxidation of NAD(P)H - the fluorescent molecule - to NAD(P)^+^. We optimized this assay to be suitable for small quantity of starting sample (60 μl) which has been mixed with 40 μl of Ammonia Assay Reagent for basal fluorescence assessment (excitation: 340 nm, emission: 460 nm). After the basal RFU acquisition, 1 μl of glutamate dehydrogenase enzyme was added to each well and, following a brief incubation, a serial acquisition of fluorescence reads was executed by Varioskan LUX (Thermo Fisher Scientific), until signal stabilization. RFU values were elaborated subtracting the last acquisition value (following enzyme addition) from the basal one. Ammonia concentration in deproteinized supernatants was calculated by using a standard curve (735 – 5880 pmol/well; LoD= 550 pmol/well) and expressed as pmol/min in relation to Seahorse assay duration. Ammonia values falling below the LoD were entered in the mathematical model as ½ * LoB (limit of blank = 6.73 pmol/min) calculated as in (23). To minimize differences with samples, all standard curve points were prepared in KREBS medium and deproteinized. Each supernatant was analysed in duplicate and the mean value was used for fluxomic calculations.

Urea concentration in deproteinized supernatants was assessed by Urea Fluorometric Assay Kit (Cayman Chemical, Ann Arbor, Michigan, USA), following manufacturer’s instructions with some modifications. The reaction volume was decreased to 100 μl and the quantity of each reagent was scaled, consistently. Specifically, 10 μl of supernatant were added to each reaction well together with 75 μl of buffer and 10 μl of enzyme (urease). After 20 min of incubation at room temperature, 5 μl of ammonia detector were added to each well and the plate was incubated for 30 min at room temperature. Then, the fluorescent signal (excitation: 410 nm, emission: 475 nm) was measured by a Varioskan LUX (Thermo Fisher Scientific). Sample concentration was calculated by using a standard curve (250 - 4000 pmol/well; LoD=250 pmol/well) and expressed as pmol/min in relation to metabolic assay timespan. To minimize differences with samples, all standard curve points were prepared in KREBS medium and deproteinized. Each sample was analysed in duplicate and the mean value was considered.

The Uric Acid Assay Kit (Merck) was performed to quantify uric acid in deproteinized media, following manufacturer’s instructions with some changes. The reaction volume was decreased to 50 μl and the quantity of each reagent was scaled, consistently. Briefly, 25 μl of samples were added to each well together with 25 μl of reaction mix (23 μl assay buffer, 1 μl enzyme mix and 1 μl probe). After 30 min of incubation, at 37°C, the fluorescent signal (excitation: 535 nm, emission: 587 nm) was measured by Varioskan LUX (Thermo Fisher Scientific). Sample concentration was calculated using a standard curve (100 - 800 pmol/well; LoD=50 pmol/well) and expressed as pmol/min in relation to assay duration. To minimize differences with samples, all standard curve points were prepared in KREBS medium and deproteinized. Each sample was analysed in duplicate. The EnzyChrom Succinate Assay Kit (BioAssays Systems, Hayward, California, USA) was used to determine succinate concentration in deproteinized media, following manufacturer’s instructions with little changes. The final volume of the reaction was adapted to 50 μl as follows: 10 μl of samples and standards plus 40 μl of working reagent (38 μl assay buffer, 0.5 μl enzyme mix, 0.5 μl cosubstrate, 0.5 μl PEP, 0.5 μl dye reagent). After 30 min of incubation at room temperature, the fluorescent signal (excitation: 530 nm, emission: 585 nm) was measured by Varioskan LUX (Thermo Fisher Scientific). Sample concentration was calculated by using a standard curve (40 – 400 pmol/well; LoD: 20 pmol/well) and expressed as pmol/min in relation to assay duration. To minimize differences with samples, all standard curve points were prepared in KREBS medium and deproteinized. Each sample was analysed in duplicate.

### Cell indirect microcalorimetry technique

The novelty of this technique relies on the transfer of indirect calorimetry principles (17, 18) to the cellular level. Classic indirect calorimetry is based on the measurement of simultaneous gaseous exchange between a bioreactor, most commonly an intact organism, and the environment (24). The rates of O_2_ consumption and CO_2_ production, together with urinary nitrogen (primarily urea) excretion, can be stoichiometrically converted into rates of glucose, lipid, protein oxidations and energy production by well-established formulae, which are valid under steady state conditions (25). The latter implies that substrate fluxes and pools are constant within the time window of study.

Carbon dioxide production rate by cultured cells is not measured in our experimental setting. However, a system of stoichiometric equations, which integrates the fluxes of four independent primary measurements - O_2_ consumption and net H^+^ release (both measured by Agilent Seahorse XFp Analyzer), lactate and NH_4_ release (both quantified by microfluorimetric methods in cell supernatants) - can substitute for the roles played by O_2_ consumption, CO_2_ production and nitrogen excretion in classical indirect calorimetry, and, hence, it can estimate (see below) the net catabolic fluxes of the three main oxidative substrates, the energy production rate, and the net fluxes/turnovers of a number of intracellular reactions/pools. Under conditions of net lipid oxidation ≥ 0, the reference metabolic map is the one in Figure 1. Please, notice that whereas substrate oxidation rates and energy production rates require only the assumptions of steady state conditions and of oxidized triglyceride/protein known compositions (see below), the estimates of fluxes/turnovers of intracellular reactions/pools require the additional assumption of no significant sub-compartmentalization of the single intracellular reactions and substrate pools.

**Figure 1.**
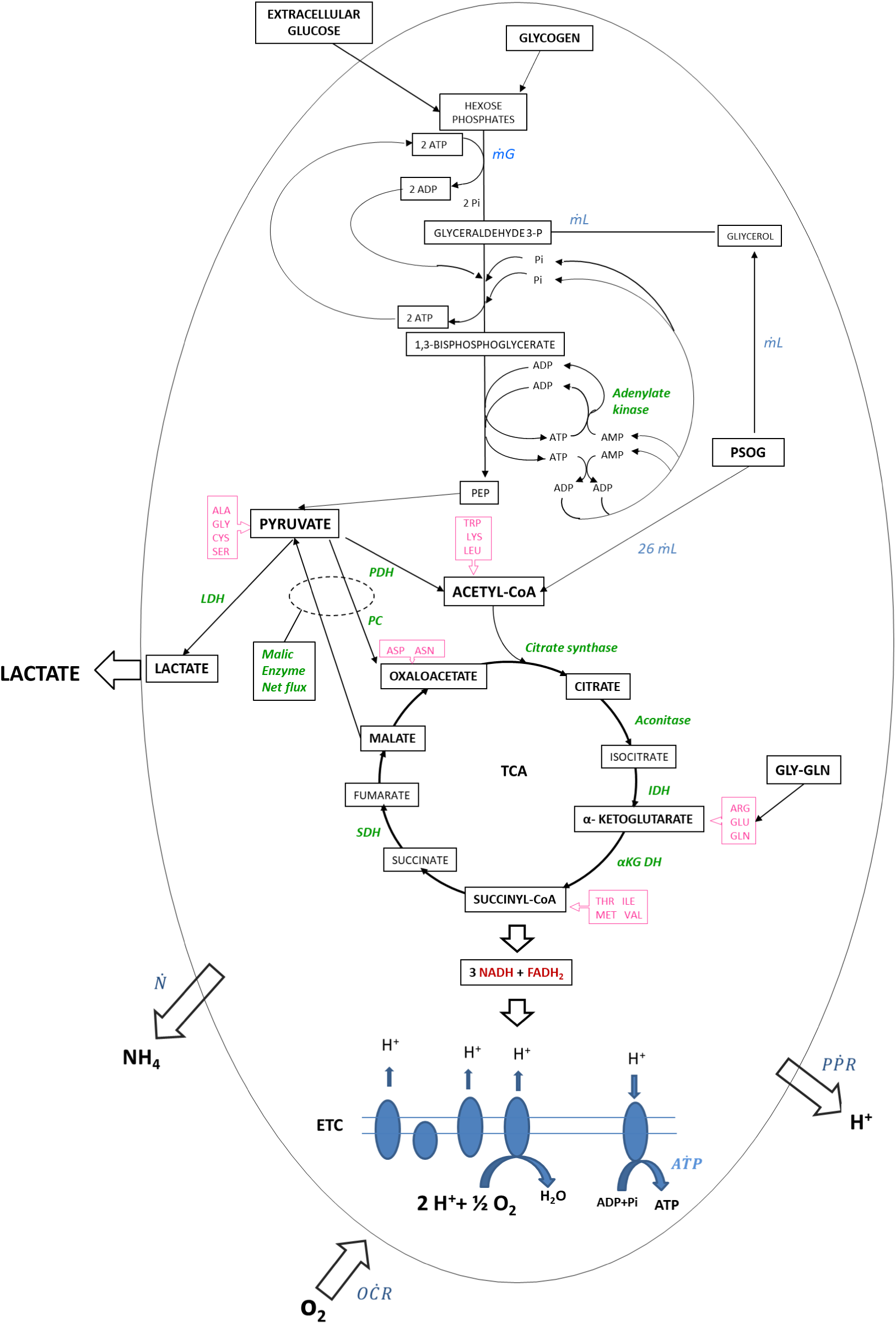
Map of the main catabolic pathways. In almost all experiments performed in this study, net lipid oxidation was ≥ 0 and this map was necessary and sufficient to account for the experimental data. Catabolic pathways are represented with black arrows (glucose, amino acids and lipids. The reference PSOG (palmityl-stearyl-oleyl glycerol) triglyceride was the reference fat for lipid catabolism. The fluxes are represented in blue. Amino acids (in pink) are shown at the points of entry into the metabolic pathways. Enzymes are named in green. (PSOG= palmityl-stearyl-oleyl glycerol, Pi= inorganic phosphate, PEP= phosphoenolpyruvate carboxylase, PDH= pyruvate dehydrogenase, LDH= lactate dehydrogenase, PC= pyruvate carboxylase, IDH= isocitrate dehydrogenase, α-KGDH= α-ketoglutarate dehydrogenase, SDH= succinate dehydrogenase, GLN= glutamine, ASP= aspartate, ASN= asparagine, VAL= valine, ILE= isoleucine, ALA= alanine, ARG= arginine, CYS= cysteine, GLY= glycine, GLU= glutamate, MET= methionine, SER= serine THR= threonine, LEU= leucine, LYS= lysine, TRP= tryptophan, ṁG = irreversible glucose utilization flux, ṁL = lipid oxidation flux, OC Ṙ= oxygen consumption flux, PP Ṙ= proton production flux, N = NH4 flux, AT Ṗ= ATP flux, ETC= electron transport chain). When the equation system of microcalorimetry yields a value of net lipid oxidation < 0, this indicates that the cell is in a mode of net lipid synthesis and a different metabolic map, and a different equation system need to be used (see Fig. S2 and Supplementary Material for the alternative metabolic map and for the modified equations).

The metabolic map of fig. 1 may strike as a gross oversimplification of intermediate catabolism, but it is built to track the net fluxes of the substrates of interest, according to the principles of indirect calorimetry. As a consequence, no futile cycle pathways are included, although their regulatory role is of recognized paramount importance (26, 27). From the viewpoint of the computation of substrate oxidation and energy production rates, under steady state conditions, including the biochemistry of futile cycles adds nothing but unnecessary complication. In the metabolic map of fig. 1, at least two cases need be discussed: the itaconate metabolism and futile cycling between pyruvate and malate/oxaloacetate.

Under steady state conditions, itaconate originates from cis-aconitate through the reaction catalyzed by IRG1. Itaconate then is converted sequentially to itaconyl-CoA and citromalyl-CoA. The latter is enzymatically split into acetyl-CoA and pyruvate (28). Pyruvate can be a substrate of pyruvate carboxylase and generate oxaloacetate, or a substrate of lactate dehydrogenase and generate lactate. In the former case, itaconate metabolism is a futile pathway embedded in the catabolism of all substrates feeding the citrate pool through pyruvate and needs not to be described in a net flux metabolic map. In the latter case, an oxaloacetate equivalent fragment of itaconate becomes part of lactate efflux from the cell, which is quantitatively described by the metabolic map, and the oxaloacetate loss is ascribed to the catabolism of one of the amino acids feeding the TCA cycle.

Also in this case, itaconate metabolism is a futile pathway embedded in the net utilization of both the amino acids feeding the TCA cycle intermediates and the substrates feeding the acetyl-CoA pool.

Being an intermediate compound of a futile pathway which needs not to be included in a metabolic map of steady state catabolic net fluxes detracts nothing from the key role played by itaconate in regulating MΦ and innate immunity, a role which depends on the size of the itaconate pool (29, 30). A similar line of reasoning applies to the reaction catalysed by pyruvate carboxylase, which generates oxaloacetate from pyruvate and CO_2_. This reaction is the reverse of malic enzyme reaction, with which it forms a futile cycle between the pyruvate and the malate/oxaloacetate pool. In the metabolic map of fig. 1, the malic enzyme flux actually represents the net balance between the unidirectional fluxes of malic enzyme and also pyruvate carboxylase, which needs not be formally included in this analysis of steady state net catabolic fluxes. Here again, this detracts nothing from the pathogenetic role of pyruvate carboxylase upregulation in monocyte derived MΦ of patients with atherosclerosis ^30^. We also retrieved our previous transcriptomic data (4) and found that, across the three subtypes of MΦ, there was no significant difference in the expression of either pyruvate carboxylase or any of the three isoforms of malic enzyme, both after 12 hours and after 24 hours of incubation with polarizing agents (data not shown).

### Substrate contribution to OĊR and PṖR

The primary parameters OĊR, PṖR, Ṅ and lactate release are expressed as pmol/min per 5×10^4^ cells, and, when entered in the appropriate equation system, enable us to compute the fluxes of interest.

The main oxidizable substrates of the cell are glucose/glycogen, triglycerides and protein/amino acids. Pyruvate is one of the primary crossroads of cellular metabolic pathways. It is primarily catabolized through two different routes: conversion to lactate by LDH or decarboxylation to acetyl-CoA. In our equation system, the fraction of net pyruvate turnover converted to lactate is defined by x, conversely 1-x represents the net pyruvate turnover fraction oxidized to acetyl-CoA. Thus, OCR (*OĊR*), PPR (*PṖR*) and ATP (AṪP) fluxes attributable to each metabolic substrate entering the pyruvate crossroad consist of three distinct components: 1. contribution of reactions before pyruvate formation, 2. contribution of pyruvate conversion to lactate (𝑥𝑥), 3. contribution of pyruvate oxidation to CO_2_ (*1-* 𝑥𝑥).

The equations of pyruvate oxidation in relation to its x and 1- x components are:

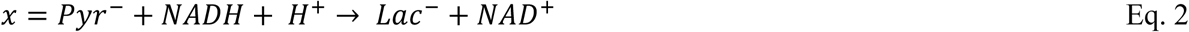

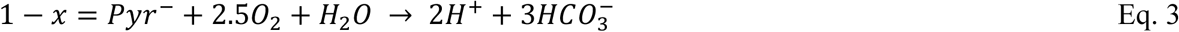

The steps and calculation used to define the equation parameters are reported below.

#### Glucose catabolism

The net equations of glucose conversion to lactate and CO_2_ are reported in eq. 4 and eq. 5, respectively:

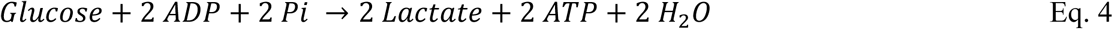

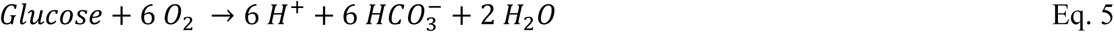

Of note, although H^+^ is generated at several sites in anaerobic glycolysis, no net H^+^ production occurs (eq. 2) unless the ATP formed is hydrolysed (12). Under steady state conditions, all ATP which is newly produced is also hydrolysed; thus, the produced ATP generates H^+^. However, ADP can be reconverted to ATP also through the phosphotransfer reaction performed by adenylate kinase and converted to adenine nucleotides (ADP and AMP) (Eq. 6), as reported in (31). This reaction (see below) has a partial buffering impact on H^+^ production rate.

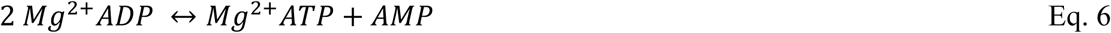

This is one example of several reactions, which in the cell can buffer partially the overall proton production rate.

To sum up, the relative contribution of glucose utilization to *OĊR* and *PṖR* expressed as a function of glucose oxidative flux (*ṁ_G_*) are described by eq. 7 and eq. 8, as follows:

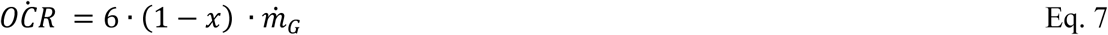

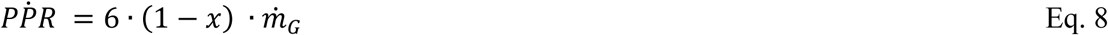

#### Amino acid oxidation

Proteins can be metabolized in different pathways, and most of the ammonium produced is released under the form of: 1. ammonium ion (NH_4_ ^+^) 2. urea (primarily in liver cells) and 3. uric acid (primary sources: liver, intestine and the vascular endothelium).

Urea and uric acid were not detected in the supernatants of MΦ in culture (data not shown). Although this finding obviously does not rule out the existence of urea and/or uric acid production by MΦ, it establishes that their quantitative contribution to the computation of amino acid oxidation is negligible. In our equation system of amino acid catabolism, therefore, we take into consideration only the pathways resulting into complete oxidation of amino acids to CO_2_, which entail simultaneous release of NH_4_.

In the absence of exogenous amino acids (no addition of glycyl-glutamine in culture media), we assumed that the cell derives the energy ascribable to amino acid from its endogenous proteins. To calculate protein oxidation parameters, we used a theoretical reference protein which contains amino acids in proportion to their abundance in the human proteome (32). We nicknamed this theoretical reference protein “proteomoid” (Eq 9). Its chemical formula is:

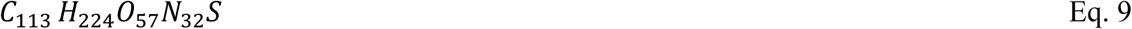

The complete oxidation of the “proteomoid” yields the following equations:

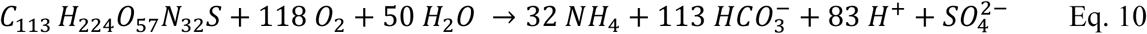

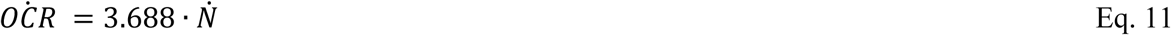

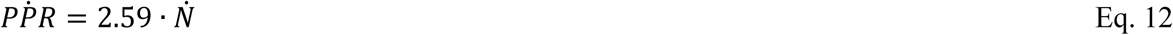

Eq. 11 and eq. 12 describe the relative contribution of complete “proteomoid” oxidation to *OĊR* and *PṖR*. However, the contribution of each amino acid to *OĊR* and *PṖR* needs be corrected for the fraction which, after reaching pyruvate, is irreversibly lost to lactate and not funnelled into the TCA cycle. Furthermore, in resemblance to many other cells, it is conceivable that macrophages may be unable to oxidize some amino acids. The “proteomoid”-derived amino acids used for irreversible catabolism, therefore, were assumed to be the sixteen amino acids listed in the supplementary table 1. Phenylalanine, tyrosine, histidine and proline were excluded from our model because their catabolism primarily occurs in the liver (33), and their metabolism in MΦ can be considered negligible for energetic purposes.

Nine amino acids directly enter the TCA cycle: arginine, glutamate and glutamine are converted to α-ketoglutarate; threonine, methionine, valine and isoleucine enter the TCA as succinyl-CoA, while aspartate and asparagine as oxaloacetate. From the standpoint of net carbon and decarboxylation fluxes, the carbons of all amino acids entering the TCA cycle downstream to the alpha-ketoglutarate dehydrogenase reaction feed the malate pool, from which, if they undergo decarboxylation, they exit the TCA cycle and enter the pyruvate pool through the malic enzyme reaction. When in the pyruvate pool, their carbons can either access the TCA cycle through the pyruvate dehydrogenase reaction or be irreversibly lost as lactate through the lactate dehydrogenase reaction. The same pattern applies to the carbons derived from glutamine, glutamate and arginine, after the decarboxylation reaction catalysed by alpha-ketoglutarate dehydrogenase.

The carbons of the other amino acids enter the TCA cycle through the acetyl-CoA pool and citrate synthase reaction. Tryptophan, lysine and leucine are converted directly to acetyl-CoA, while the carbons derived from alanine, glycine, cysteine and serine end up in the pyruvate pool, and undergo the metabolic fates described above (Fig. 1).

In several experiments, we endeavoured to measure succinate release in the medium under the two polar conditions of no substrates and GlyGln plus glucose in the culture medium. Succinate was selected because of its unique position of being a reflex of the reaction catalysed by SDH, which in turn is located in the unique position of belonging to both the TCA cycle and the electron transport chain pathway. A loss of succinate by the cell should be accounted for in the stoichiometric equations of both the TCA cycle and the aerobic ATP production pathway. Our determinations of succinate were below the detection limit of the assay. Thus, in our equations we did not include any succinate exit from the TCA cycle.

For all amino acids which contribute to the pyruvate pool, the x and 1- x components were included in the stoichiometric equations of the model (see the equations in the Supplementary Table 1).

The equations of amino acid oxidation to CO_2_ and the relative fluxes of *OĊR* and *PṖR* are expressed as a function of ammonia fluxes (*Ṅ*) (Supplementary Table 1).

When glycine-glutamine was added to the culture medium, NH_4_ release could be originated by either the “proteomoid” or Gly-Gln. We assumed that Gly-Gln would be predominant and the ratio GlyGln/”proteomoid” of amino acid derived carbon influx into the pyruvate and α-ketoglutarate pools would be 3:1.

#### Lipid oxidation

In order to determine endogenous lipid contribution to cell bioenergetics, we selected, as in indirect calorimetry (17, 18), palmitoyl-stearyl-oleyl-glycerol (PSOG) as the reference triglyceride. The oxidation of PSOG to pyruvate and CO_2_ yields to the following equations:

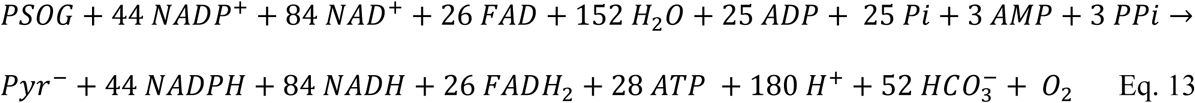

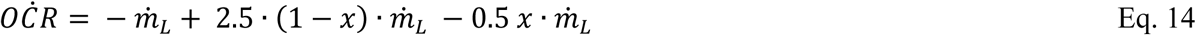

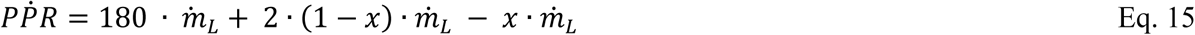

in which, equations 14 and 15 represented the relative contribution of PSOG oxidation to *OĊR* and *PṖR* expressed as a function of lipid oxidation flux (*ṁ_L_*). Please note that, under steady state conditions, the process goes on by funnelling, directly or indirectly, NADPH, NADH, and FADH_2_ into the oxidative phosphorylation pathway, in which O_2_ consumption takes place.

In a little number of cases (n=6; 2 in M0, 3 in M1, and 1 in M2 MΦ) the equations of lipid oxidation yielded negative fluxes, i.e. net lipid oxidation < 0. They occurred with substrate deprived medium (n=4) or with 5 mM glucose added medium (n=2). In indirect calorimetry studies, negative lipid oxidation proofs that the cell is in a metabolic mode of net fat synthesis (17, 18). In such cases, in analogy to indirect calorimetry, we used a different metabolic map and different equations to compute net fat (PSOG) synthesis and to provide correct estimates of net glucose/amino acid oxidation. The modified metabolic map and equation system are reported in the Supplementary Material.

### Energetic Fluxes

Energy turnover was computed exploiting the standard enthalpies of combustion of substrates and the standard enthalpies of formation of products, according to the principles of indirect calorimetry. Slightly differences in the equations are due to the substrate admixture which is assumed to be oxidized, i.e. glycogen-PSOG-“proteomoid” or glucose-PSOG-“proteomoid” and so on. For instance, in the case of the culture medium with no substrates, the admixture is glycogen-PSOG- “proteomoid”. Under this experimental setting the equation for energy production rate (EPR) is:

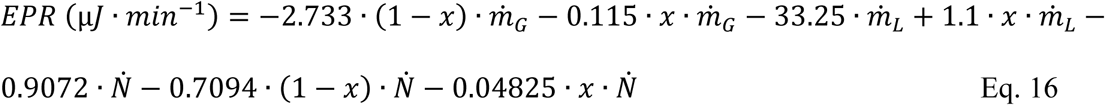

EPR equations are written with the minus sign, because this energy output also is an energy “loss”. The other equations for EPR can be found in the Supplementary Material.

Please, note that, as long as the primary measures are correct, the computation of EPR is very reliable, being firmly based on thermodynamics.

On the basis of known biochemistry pathways, we also computed maximum, aerobic and anaerobic ATP turnover (pmol/min) (see equations in the Supplementary Material). Please, notice that the computation of aerobic ATP synthesis assumes complete coupling between the generation of reducing potential and the production of ATP. Since we have measured no biomarker of oxidative phosphorylation coupling, aerobic and maximum ATP synthesis are upper boundaries of numbers which remain unknown. On the other hand, cell indirect microcalorimetry EPR, like indirect calorimetry (17, 18), does not suffer from this uncertainty.

### Metabolic flux analysis

As already mentioned, the net stoichiometry of the oxidative pathways results into a system of equations (Supplementary Material), based on four measured primary fluxes (OCR, PPR, lactate and NH_4_) used to derive four primary outputs: net irreversible glucose utilization rate (*ṁ_G_*), net lipid oxidation (*ṁ_L_*), pyruvate dehydrogenase flux (PDH), the fraction of H^+^ (*ω*) which escapes the intracellular buffering mechanisms, including e.g. adenylate kinase reaction. Please note that the primary measure of lactate release in the medium provides the rate of pyruvate conversion to lactate.

The rate of amino acid oxidation is primarily derived from NH_4_ data coupled with the known “proteomoid” composition ± the glycyl-glutamine contribution. Please, note that by this approach the admixture of amino acids irreversibly oxidized by the cells is assumed to be known, thereby opening a window on several pathways.

Twenty-four other unknown fluxes/parameters can be computed downstream to these primary determinations. For each of them, slightly different equations are needed as a consequence of the substrates present/absent in the culture media (e.g. if glucose is absent in the medium we assume that only glycogenolysis is the energetic pathway contributing to glucose oxidation; if no glycyl-glutamine is added, only the “proteomoid” is the source of amino acid oxidation and NH_4_ production, and so on).

The model with the primary measurements of each experiment was implemented in the modelling software SAAM II (34), in which identification of the four primary unknown parameters (*ṁ_G_*, *ṁ_L_*, *PDH*, and ω) and computation of all other derived fluxes and metabolic parameters were carried out (Supplementary figure S1).

Please, note that all fluxes, in agreement with the indirect calorimetry foundations on thermodynamics, are net fluxes. The fluxomic parameters can be divided into three categories:

1. Substrate fluxes: Irreversible Glucose Utilization, Net Lipid Oxidation, Irreversible Amino Acid Utilization, Glucose Oxidation, Anaerobic Glycolysis;
2. Oxidative fluxes: Pyruvate Input, pyruvate dehydrogenase (*PDH*), Aconitase, α-ketoglutarate dehydrogenase (aKGDH), succinate dehydrogenase (SDH), TCA Net Flux, Citrate Net Synthesis, Malic Enzyme, Malate Turnover;
3. Bioenergetic fluxes/ratios: Energy Production Rate/Expenditure (EPR), Anaerobic EPR, Max ATP production, Anaerobic ATP, Energy Production Efficiency, Aerobic Energy Production Efficiency, P/O ratio, Aerobic P/O ratio.

Energy substrate fluxes of the catabolic pathways are expressed in pmol/min, and include lipid oxidation, irreversible amino acid utilization (oxidation), and irreversible utilization of glucose, which is the sum of anaerobic glycolysis (i.e. most of lactate production) and glucose oxidation. The pivotal oxidative flux is the net TCA flux (pmol/min), which is expressed as the flux of all acetyl-CoA equivalents turning over the cycle. Other oxidative fluxes are listed above.

Bioenergetic fluxes include fluxes of maximum and anaerobic ATP turnover (pmol/min) and energy production rate/expenditure (expressed in μJ/min). Maximum ATP turnover is a flux which is computed assuming no uncoupling of the oxidative phosphorylation, hence it provides the upper bond of true total ATP turnover.

The efficiency of energy production is also computed. The latter has no units, i.e. it is a pure number, because it is the ratio between the energy “packed” in total ATP production rate (converted in μJ/min) and the sum of total energy production rate/expenditure (i.e. heat, μJ/min) plus lactate outflux (mass flux, i.e. pmol/min, expressed in μJ/min equivalents) (eq. 17).

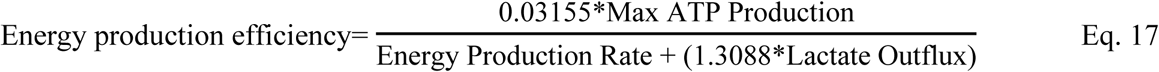

## Statistical analysis

All fluxes are normalized to 5 x10^4^ cells. All data are presented as mean±SEM. Comparisons between two different culture conditions were carried out by paired t-test, whilst repeated measure ANOVA (followed by Dunnet post-hoc) or Friedman (with Dunn’s post-hoc) tests were used to detect differences among the different culture media. Differences among M0, M(LPS+INF-γ) and M(IL-4) were investigated by one-way ANOVA with Tukey post-hoc. Statistical significance was set at p<0.05 (two-sided). Data analysis was performed using SPSS version 24.0 (SPSS Inc/IBM, Chicago, Illinois, USA) and GraphPad PRISM version 10.0.2 (GraphPad Software, Boston, Massachusetts, USA).

## Results

### Influence of substrate supply on the bioenergetic fluxes of human macrophages

The implementation of the metabolic equations in the SAAM II software and of the four primary measures of each experiment (lactate release, NH_4_ release, OCR and PPR) provided us with the numerical identification of the four unknown parameters (irreversible glucose utilization, lipid oxidation, PDH flux, and *w*, the fraction of H^+^ which escapes the acid buffering capacity of the cell) (Supplementary figure S1) and the computation of all other metabolic fluxes, listed in the stoichiometric equations.

As a preliminary test of the experimental system sensitivity to metabolic changes, the MΦ metabolic fluxes were quantified in KREBS medium under four different conditions of substrate availability: i. absence of exogenous substrates, ii. glucose 5 mM, iii. glycyl-glutamine 2 mM, iv. glucose 5 mM + glycyl-glutamine 2 mM.

All primary measures as well as the derived fluxomic parameters are presented in table 1. The main differences in primary measures were found in lactate production rate, whereas changes in O_2_ were hardly detectable.

**Table 1.**
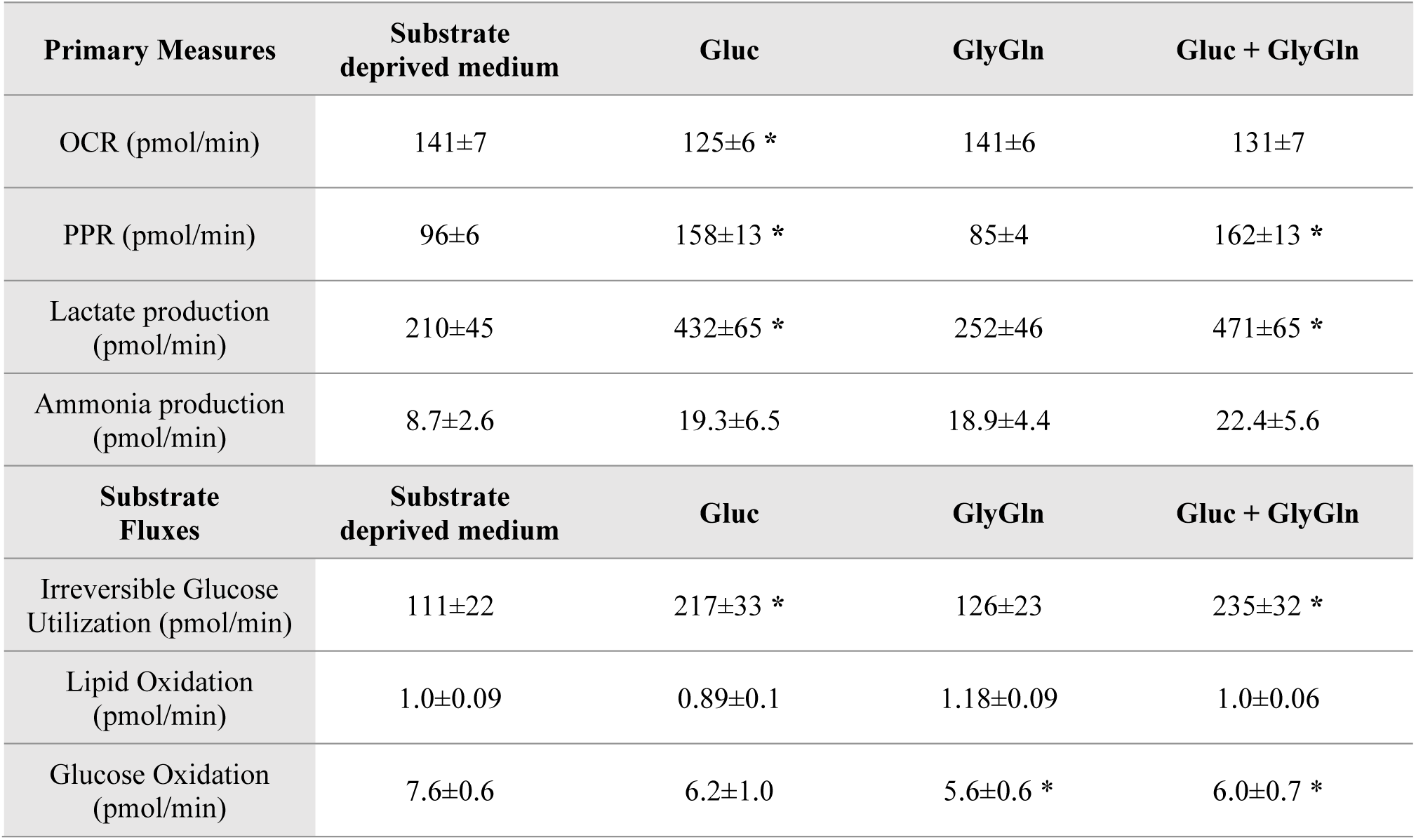
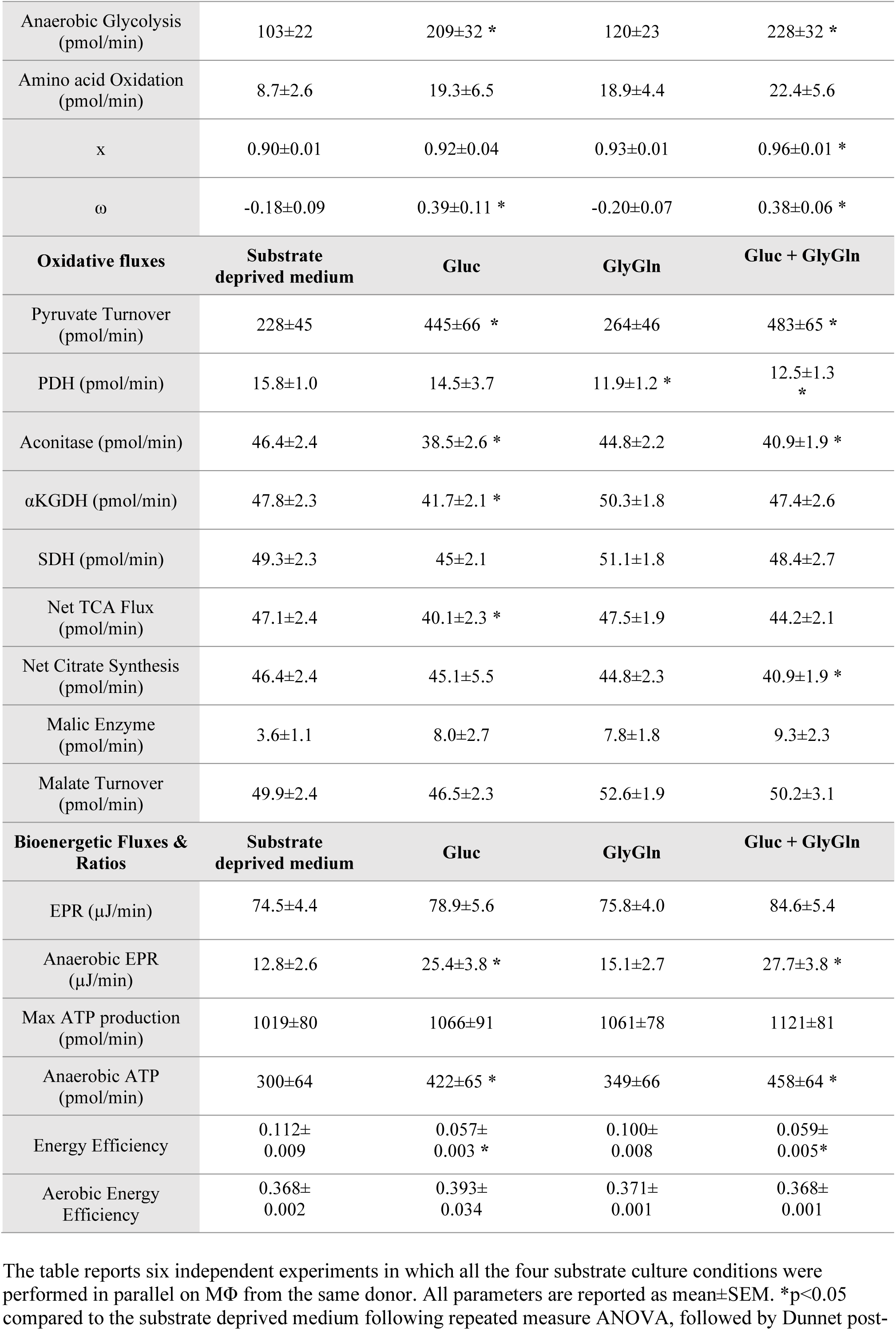
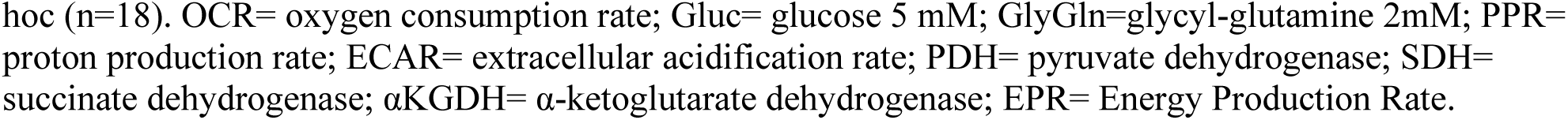
Fluxomic parameters of primary human macrophages exposed to different metabolic substrates.

When proton production rates and lactate outflux rates were plotted against each other (figure 2), the relationship, albeit statistically significant, was weak and far from being 1:1. Furthermore, in the presence of glycyl-glutamine, the association was weaker, when compared to no exogenous substrates or to glucose alone.

**Figure 2.**
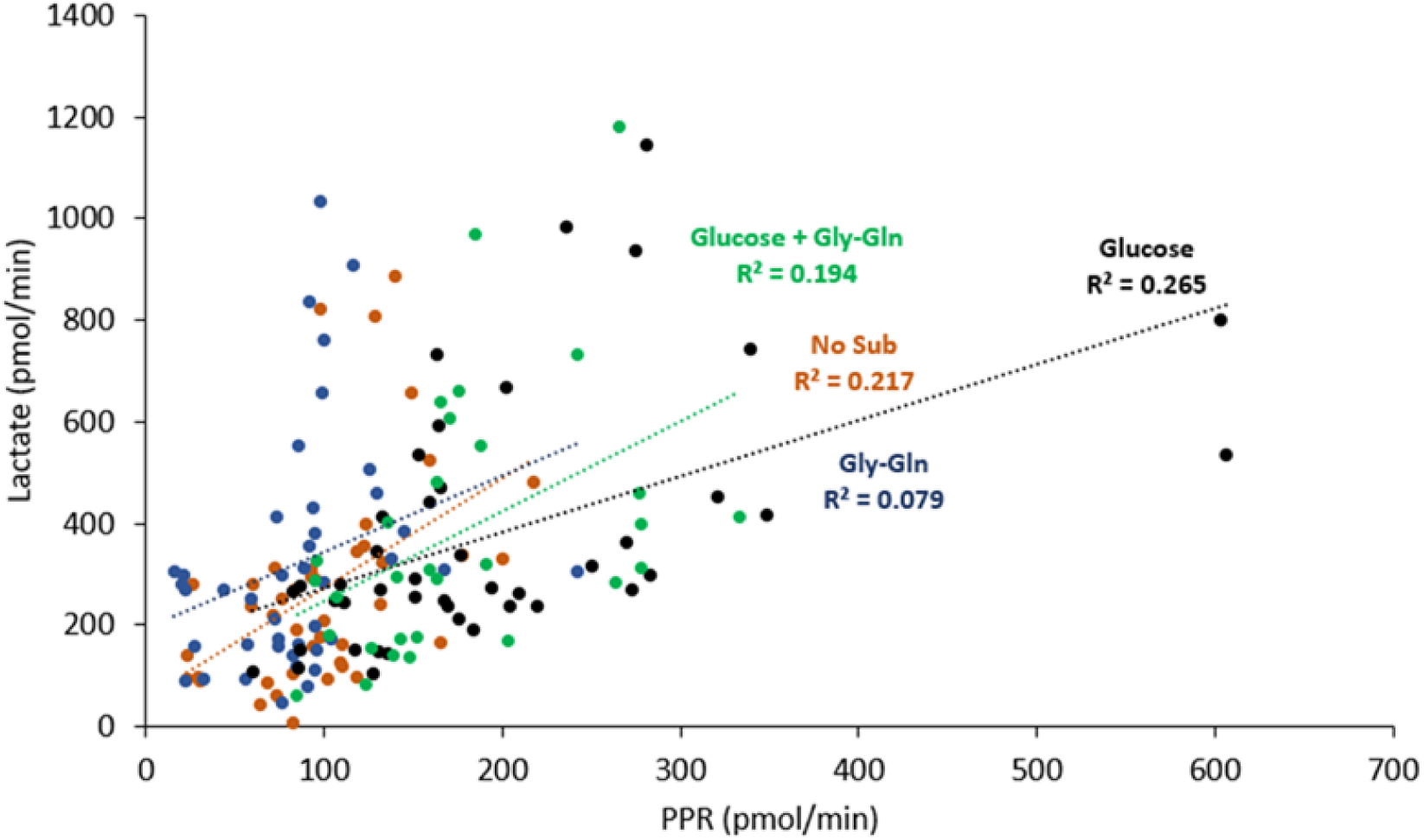
Correlation between lactate and PPR levels. Correlation between the primary measures of lactate (by enzymatic assay) and proton (by Seahorse) release rates of M0, M(LPS+INF-γ) and M(IL-4) exposed to different media (with no additional substrate, glucose 5mM, glycyl-glutamine 2mM, or glucose + glycyl-glutamine) during metabolic assay.

When glucose was present in the medium, there was a striking increase in the irreversible glucose utilization rate, entirely accountable for by anaerobic glycolysis and lactate production, whereas no increase in glucose oxidation through PDH was detected. With glucose alone, TCA cycle somewhat slowed down, as shown by a decrease in net TCA cycle flux, aconitase flux and aKGDH flux. The efficiency of energy production fell because of the increase in lactate release, which, from the bioenergetic viewpoint, is an irreversible loss of energy equivalents.

When GlyGln was present in the medium, there was a small, but significant, decrease in glucose oxidation and in PDH flux.

With glucose alone and GlyGln alone, there was a similar, but not significant, increase in x, which quantifies the ratio of LDH flux divided by the sum of LDH pus PDH fluxes. When both exogenous sources (glucose and glycine) of pyruvate where present in the culture medium, x increased further and reached the statistical significance in the comparison to the no substrate condition.

When glucose was present in the culture medium, omega increased significantly, i.e. less protons were buffered by intracellular reactions.

Throughout the four conditions, there was a remarkable stability in EPR, maximum ATP production rate and efficiency of aerobic energy production.

### Macrophage phenotypic and functional characterization

*In vitro* polarized human monocyte-derived macrophages were investigated for M1 and M2 signatures. M(IL-4) displayed higher expression levels of the M2-specific surface markers (35–37). CD200R and CD209 compared to both M0 (unstimulated) and M(LPS+INF-γ). CD206 was significantly increased only in M(IL-4) compared to M(LPS+INF-γ). Moreover, M(LPS+INF-γ) showed higher levels of the surface markers CD80 (Figure 3A) compared to both M0 and M(IL-4), as expected (38, 39). The expression of markers CD80 and CD209 was confirmed by flow cytometry, showing an increased CD80 and CD209 expression in M(LPS+INF-γ) and M(IL-4), respectively (Figure 3B). As for macrophage function, we found that M(LPS+INF-γ) showed a higher expression of the pro-inflammatory markers IL-1β, TNF-α, IL-8, IL-6 compared to both M0 and M(IL-4) (Figure 3C), documenting their pro-inflammatory mode.

**Figure 3.**
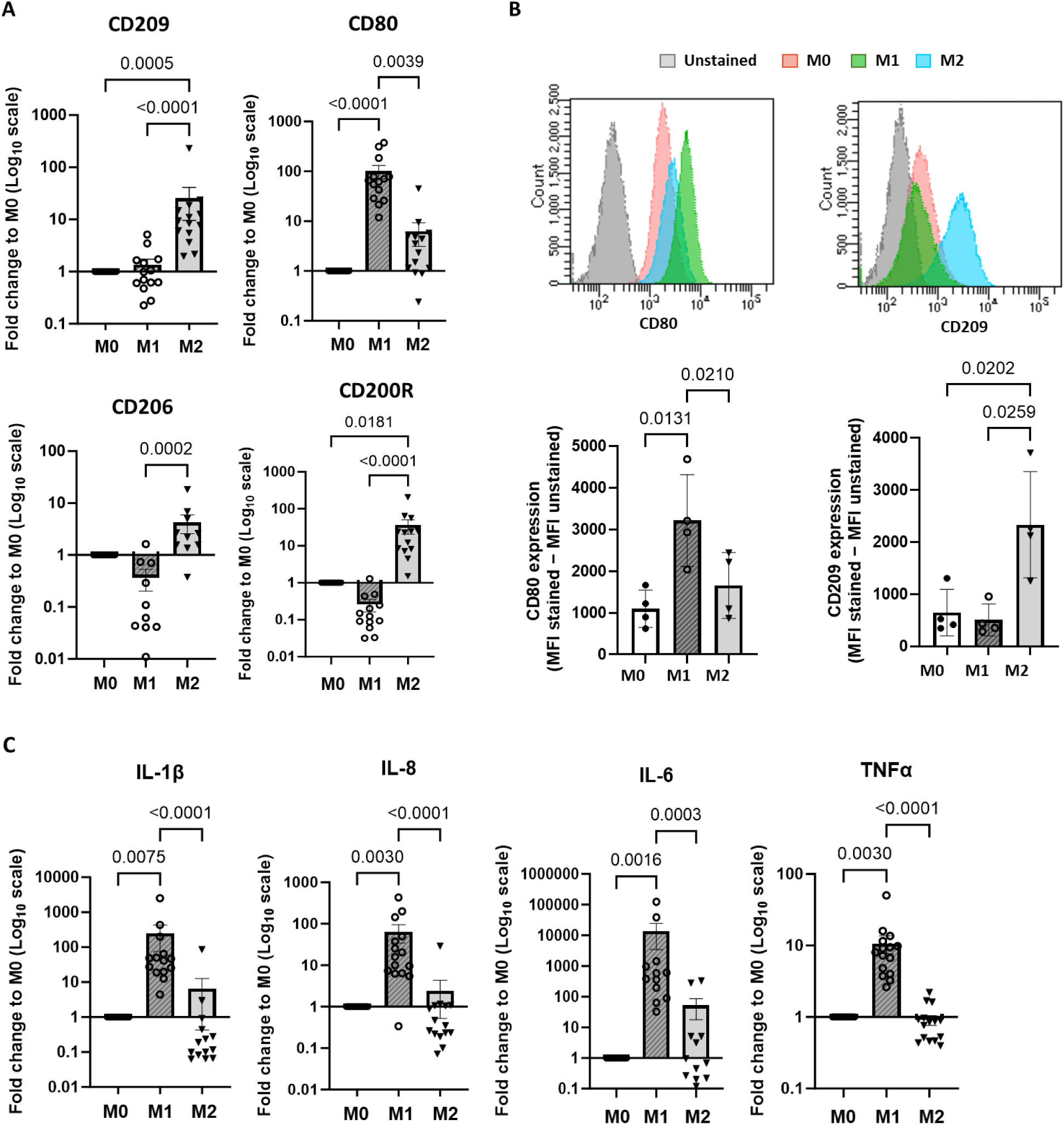
Identity of polarized human monocyte-derived macrophages. To evaluate the polarized status of macrophages, the gene expression of selected cell surface markers (CD80, CD200R, CD209, CD206) **(A)** and pro-inflammatory genes (IL-1β, TNF-α, IL-8, IL-6) **(C)** was evaluated by qPCR. The expression of CD80 and CD209 was confirmed by FACS analysis **(B)** in which MFI values were corrected for autofluorescence by subtracting the mean fluorescence intensity (MFI) of the unstained control from the MFI of the stained sample (i.e., ΔMFI = MFI stained – MFI unstained). Data are expressed as mean±SEM and analysed with Friedman test. IL= interleukin; TNF= tumor necrosis factor; M0=unstimulated macrophages; M1= M(LPS+INF-γ); M2= M(IL-4).

### Metabolic characterization of M(LPS+INF-γ) and M(IL-4) macrophages

The bioenergetic metabolic fluxes assessed by indirect microcalorimetry in each culture medium were analysed further by the three subtypes of MΦ (M0, M(LPS+INF-γ) and M(IL-4)). All primary measures and derived metabolic fluxes are presented in table 2. Importantly, no significant changes in proton leak rates were found across the three MΦ subtypes (Table S4).

**Table 2.**
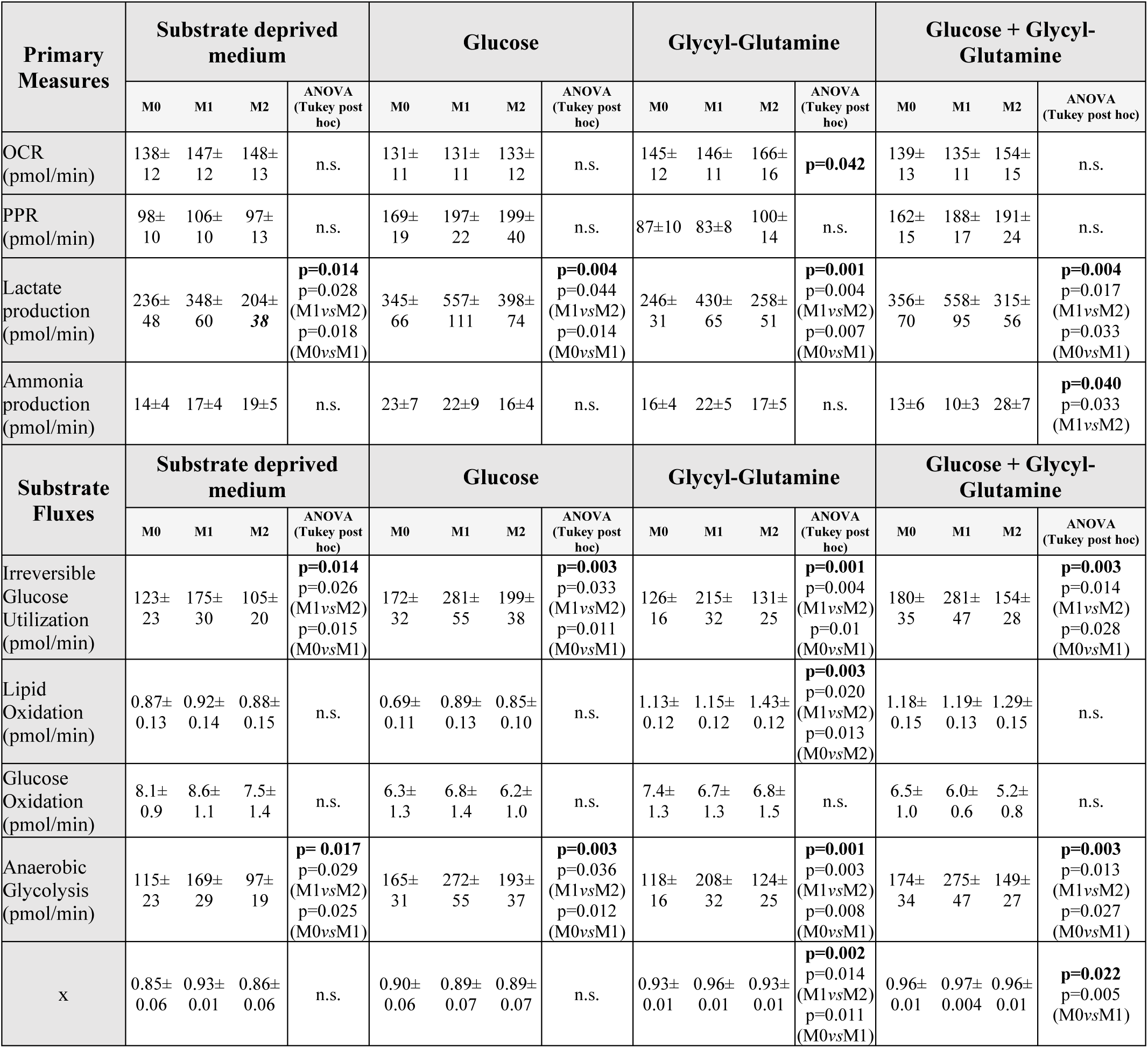
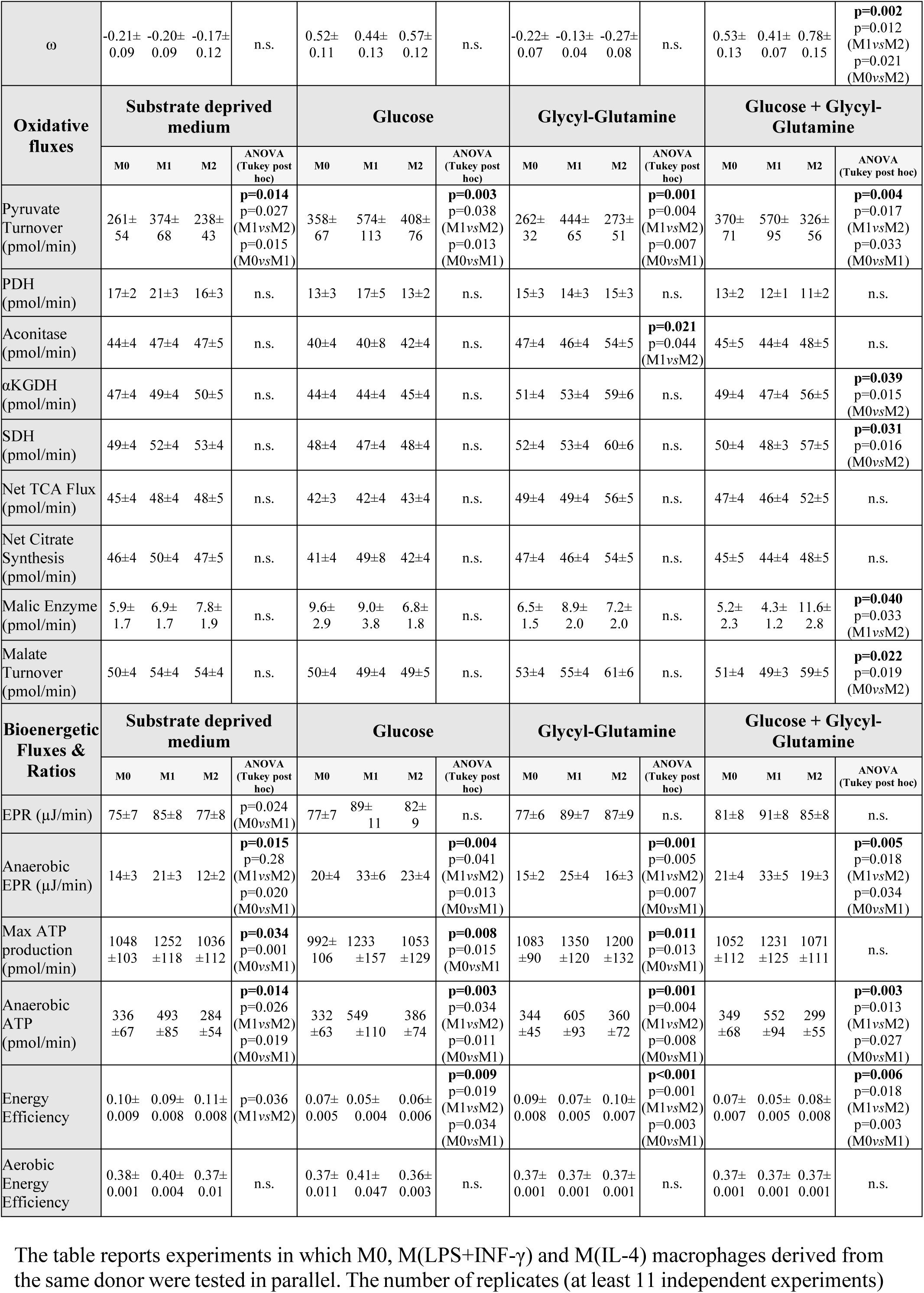
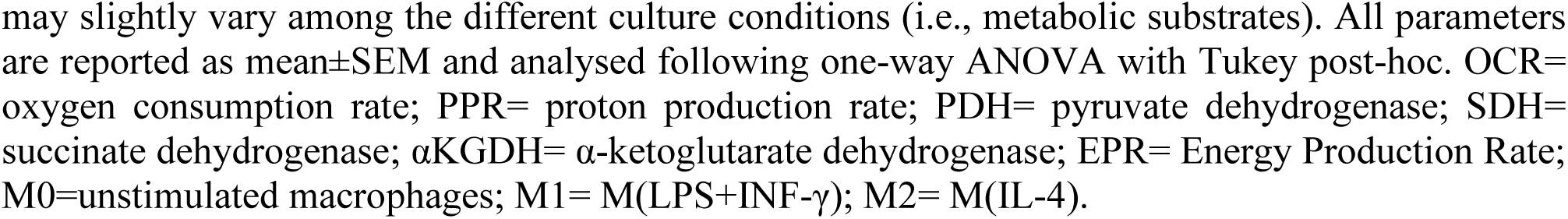
Fluxomic parameters of M0, M(LPS+INF-γ) and M(IL-4) monocyte-derived macrophages exposed to different energetic substrates.

There was a trend to higher oxygen consumption in M(IL-4) when exposed to GlyGln supplemented media. Lactate release was significantly greater in M(LPS+INF-γ), across all four media, whereas PPR showed no difference at all. Finally, ammonia release was significantly greater in M(IL-4) than in M(LPS+INF-γ) in glucose + GlyGln supplemented medium.

Glucose oxidation and PDH flux were similar in all three macrophage subtypes, irrespective of culture medium. In contrast, irreversible glucose usage, anaerobic glycolysis and pyruvate turnover were significantly higher in M(LPS+INF-γ) than in M(IL-4) or M0. Lipid oxidation was higher in M(IL-4) than in M(LPS+INF-γ) or M0 only in GlyGln supplemented medium.

Although net TCA flux showed only nonsignificant trends, aKGDH and SDH fluxes were significantly higher in M(IL-4) than in M(LPS+INF-γ) or M0 in glucose + GlyGln supplemented medium. In this same medium malic enzyme flux was higher in M(IL-4) than in M(LPS+INF-γ).

Energy production rates were similar across the three macrophage subtypes, irrespective of culture media. However, anaerobic EPR, maximum and anaerobic ATP production were consistently higher in M(LPS+INF-γ) than in M(IL-4) or M0. In contrast, energy efficiency was consistently lower in M(LPS+INF-γ) than in M(IL-4) or M0. Aerobic energy efficiency was similar across the three macrophage subtypes, irrespective of culture medium.

In figure 4, all measured and derived metabolic fluxes of M(LPS+INF-γ) and M(IL-4) in glucose + GlyGln supplemented medium are presented to highlight the fluxomic signatures of these two macrophage subtypes. M(LPS+INF-γ) featured accelerated anaerobic glucose metabolism and anaerobic ATP production, at the expense of lower energy production efficiency. M(IL-4) displayed a two-speed TCA cycle: not significantly higher than M(LPS+INF-γ) between malate and a-ketoglutarate, but faster than M(LPS+INF-γ) in the route from a-ketoglutarate to malate, with higher malic enzyme flux to pyruvate, which enables malate dehydrogenase and citrate synthase to somewhat slow down, close to the rate seen in M(LPS+INF-γ).

**Figure 4.**
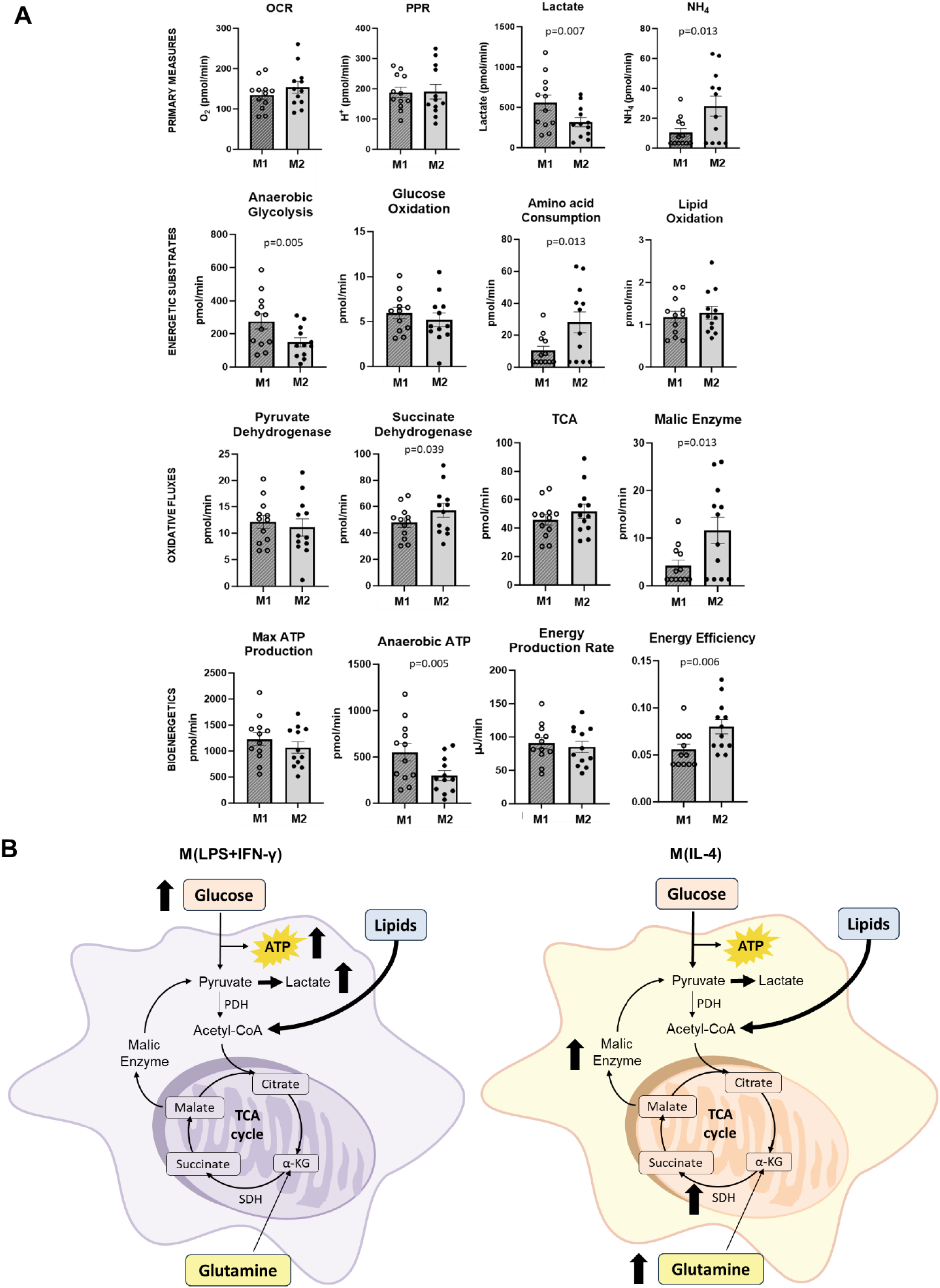
Fluxomic parameters of M(LPS+IFN-γ) and M(IL-4) macrophages cultured with glucose and glycyl-glutamine. Panel A shows the main fluxomic parameters (pmol/min) of M(LPS+IFN-γ) (indicated as M1, closed circles) compared to M(IL-4) (indicated as M2, filled circles) under culture conditions containing 5 mM glucose and 2 mM glycyl-glutamine. Data are reported as mean±SEM of at least 10 independent experiments following paired t-test, only p<0.05 are shown. Panel B provides a schematic overview of the differences in metabolic fluxes. OCR= oxygen consumption rate; PPR= proton production rate; TCA= tricarboxylic acid cycle.

## Discussion

In this paper, we have introduced a novel technique of cell indirect microcalorimetry which enables to estimate cell metabolic fluxes, and, by exploiting this approach, we have outlined basal bioenergetic metabolism of primary human MΦ after exposure to different cell polarizing media.

Thanks to the acquisition of four primary measures (i.e. OCR, PPR, lactate and NH4 releases), we can compute, in a time-and cost-efficient way, the main parameters related to cellular metabolic pathways under (quasi-)steady state conditions, with a special focus on lipid, glucose and amino acid oxidation, anaerobic glycolysis, tricarboxylic acid cycle (TCA), energy expenditure and efficiency, maximum and anaerobic ATP production.

Nowadays, most research laboratories assess oxygen consumption rate (OCR) and (extra-cellular acidification rate) ECAR or proton production rate (PPR) to investigate cellular bioenergetic metabolism (40). Previously published papers have greatly improved the correct interpretation of the data provided by this approach and introduced methods to correct for some of its limitations (15).

In this study, we have endeavoured to further progress on the path of better clarifying the meaning and the use of the primary data (OCR and PPR) regarding the general problem of cellular bioenergetic metabolism. We were inspired by the principles of indirect calorimetry (18), which in turn are firmly based on thermodynamics (17). From this standpoint, under our experimental conditions it was mandatory:

1. To introduce a readout of protein/amino acid oxidation
2. To select an intracellular “reference protein” available for oxidation by the cell
3. To take into account CO2 production, in the absence of a direct measure
4. To experimentally test the assumption that PPR, as derived from ECAR, is a close readout of lactate production.

The first and second point are obviously linked to each other. The third and the fourth point also are intertwined, because CO_2_ production also is an acidifying process, which, for instance, fully manifests itself in human pathology with the respiratory acidosis due to respiratory failure (41).

In human primary MΦ we found that the release of uric acid and urea was undetectable, thereby leaving NH_4_ as the only detectable readout of amino acid oxidation. We then selected a reference virtual protein (“proteomoid”) with the same amino acid composition of human proteome (32) and derived the stoichiometric equations of its catabolism. This also is the strategy used in whole body indirect calorimetry in humans (42). In the absence of amino acids in the medium and given the minimal or absent *in vitro* production of NO of human MΦ (43), which incidentally confers our species vulnerability to Rickettsial infections (44), the NH_4_ release is informative of irreversible amino acid catabolism.

At the same time, our data show that in human primary MΦ the relationship between PPR and lactate release is clearly too weak to enable the former to be a reliable readout of the latter (Fig. 2). We therefore argued that the PPR reflects more than just lactate release in the medium, and it is the net result of several processes, including anaerobic glycolysis, CO_2_ production, amino acid catabolism, proton buffering reactions, such as those catalysed by adenylate kinases, etc.

All these considerations were crystallized in sets of stoichiometric equations (see also Supplementary Material), which can be used to quantify the net fluxes through the main cell bioenergetic pathways based on the four primary measures assessed in this study.

We first tested cell indirect microcalorimetry to portray fluxomics of human monocyte-derived MΦ in the presence/absence of two metabolic substrates, representative of glucose and amino acid (glycyl-glutamine) metabolism, with the idea of testing whether this experimental system was sensitive enough to detect metabolic responses to the changes caused by exogenous substrate availability. Please, note that, since GlyGln supplies only two amino acids, in all experimental conditions herein reported the MΦ were in protein catabolic state. Thus, as a net balance, GlyGln can contribute only to amino acid oxidation.

This study shows that, in close parallel to other studies (45), human MΦ release a large amount of lactate, with figures between 13 and 27 nmoles per hour per 5 x 10^4^ cells, highly comparable to the rates reported in (45).

Proton efflux is widely considered a readout of anaerobic glycolysis and is assessed often in the place of lactate (15, 40). However, only a weak and lower than a 1:1 correlation between PPR and lactate release was found in our study, particularly in the absence of exogenous glucose (fig. 2).

This finding proves that in human MΦ PPR does not mirror anaerobic glycolysis, but it is the net result of several proton producing and proton buffering reactions, as already suggested (46). Thus, the direct measure of lactate release should be considered a key requirement in the assessment of human MΦ metabolism. A corollary of this finding is that, at least in human MΦ, the widespread habit of using PPR as a proxy of anaerobic glycolysis may be misleading (47–50).

As to the primary measures, in our study OCR did not change across the different culture conditions, while in the presence of glucose in the medium lactate release rate approximately doubled and PPR increased by 50-70%. Although highly variable, NH_4_ release rate, our readout of amino acid catabolism, did not change significantly across the different substrate supplies.

In the presence of glucose 5 mM, the increases in irreversible glucose utilization and anaerobic glycolysis, coupled with the absence of changes in glucose oxidation, demonstrate that under these experimental conditions glucose transport and the reactions catalysed by hexokinase and PFK are not rate limiting in human MΦ, in spite of the absence of stimuli of glucose transport/phosphorylation, such as insulin (51). In contrast PDH appears to be a rate limiting step, which strictly regulates the access of glucose/glycogen derived carbons to the TCA cycle, as shown by the tiny fraction of glucose/glycogen usage undergoing oxidation (2.6 – 6.8%). Nevertheless, the contribution of glucose derived acetyl-CoA to net citrate synthesis was rather stable around 28-33% across the different substrate conditions, again highlighting the role played by PDH also in the regulation of the TCA cycle (52). Conversely, the lipid share, easily computed by multiplying net lipid oxidation by a factor of 27, accounted for 53-71% of net citrate synthesis, being the major contributor under all the metabolic conditions herein explored.

Thus, apart from anaerobic glucose metabolism, the fluxomic signatures of human primary MΦ appear to be relatively insensitive to acute changes of substrate supply *per se*.

Glucose supply in the medium and the concomitant acceleration in anaerobic glycolysis, on the other hand, overcome the capacity of proton buffering reactions, as reflected by the significant increase in the parameter w. Although the rate of proton release increases, its relationship to the rate of anaerobic glycolysis remains quite weak (fig. 2). The variable efficiency of intracellular buffering reaction(s) is an additional reason for not relying on PPR to track anaerobic glycolysis.

Energy production rate (EPR value) of MΦ was about 70-85 µJ/min, a lower figure than the value (96 µJ/min_)_ of energy expenditure found in a previous study with direct calorimetry.

However, in that study murine, not human, MΦ were investigated (53). It is well known that hypometric scaling underlies the change of EPR across species, i.e. species with larger bodies have lower EPR per mass unit than small body species (54), a finding tentatively ascribed also to lower proton leak across the mitochondrial membrane in large body species (55). The latter explanation would affect also EPR in cultured mammalian cells and would predict that MΦ EPR in humans should be 80-90% of the figure found in mice. Our findings are compatible with these predictions (55). The absence of substrate driven significant changes in EPR (table 1) is consistent with the general rule of a tight regulation of TCA cycle/ETC pathways (52), which accelerate only when uncoupled or under conditions of increased energy demand by the cell.

When the data were analysed by layering them according to the polarization protocol, we confirmed that the exposure of donors’ monocytes to cell polarizing agents was successful in achieving clearcut different phenotypes. MΦ exposed to LPS+IFN-g expressed the surface marker CD80, whereas MΦ exposed to IL-4 expressed the surface markers CD200R, CD209 and CD206. The former and the latter pattern hallmark successful M(LPS+INF-γ) (former M1) (35, 38, 39) and M(IL-4) (former M2) polarization (35–37, 56), respectively. Furthermore, the differential expression of a set of inflammatory markers (IL-1β, TNF-α, IL-8, IL-6) are further evidence that we generated typical M(LPS+IFN-g) and M(IL-4) polarized phenotypes.

The cell indirect microcalorimetry technique herein introduced enabled us to outline the metabolic features of human polarized MΦ. Please note that the absence of significant changes in proton leak between the three MΦ subtypes is compatible with the presence of similar coupling between OCR and ATP production across the three MΦ subtypes.

Under no condition PPR was different across the three MΦ subtypes, further demonstrating that PPR may be an insufficient readout to detect immune-metabolic differences across human MΦ subtypes and to track glucose/glycogen catabolism. However, the anaerobic glycolytic flux of M(LPS+INF-γ) was about twice the figures M0 and M(IL-4) under all conditions, including the absence of substrates in the culture medium. These data align the metabolic behaviour of human primary MΦ to a huge number of studies in rodent MΦ and confirm the presence of Warburg effect in human primary M(LPS+INF-γ) (8, 9).

Quantitation of TCA cycle by cell indirect microcalorimetry was consistent with a rather preserved cycle in M(LPS+INF-γ), a finding which apparently is at variance with some published evidence (8). It is well established that LPS-stimulated murine MΦ exhibit a dual alteration of the TCA cycle, characterized by both succinate accumulation and itaconate production (49, 57). However, at normoxic conditions, M(LPS) show no increase in reductive carboxylation of a-ketoglutarate and unchanged channelling of glutamine derived carbons from a-ketoglutarate to malate, both findings pointing to intact clockwise turning of the TCA cycle (58). Furthermore, our results are similar to studies which compared human and murine MΦ and reported that human and mouse MΦ have different aerobic metabolism in response to LPS (45, 59).

In our studies, when exposed to no, or just one, substrate, human M(LPS+INF-γ) revealed no changes in OCR and TCA fluxes compared to the M(IL-4) population, as a further indication of an efficient TCA cycle. This does not contradict the increases in intracellular succinate and itaconate reported also in human pro-inflammatory monocyte derived MΦ (9): fluxes may stay the same at steady state in spite of variations in substrate levels, which in turn hallmark changes in the catalytic activity of the enzymes. Indeed, flux rates and quantitative metabolomics are complementary to each other.

The simultaneous exposure of MΦ to both glucose and GlyGln was more instructive in unveiling differences between MΦ subtypes. M(IL-4) confirmed to have a high oxidative profile and increased TCA-related fluxes compared to M0 (table 2). It should be noted that the acceleration of TCA cycle in M(IL-4), when cultured in glucose + GlyGln medium was seen only downstream to aconitase, i.e. at the site of entry of glutamine derived a-ketoglutarate. M(IL-4) had also a higher malic enzyme flux than M(LPS+INF-γ), thereby funnelling more malate into the pyruvate pool and resulting into a citrate synthase flux which was close to the rates seen in M(LPS+INF-γ) and M0.

This pattern further suggests that PDH plays a key role in controlling the rate of the TCA cycle in human primary MΦ.

On the other hand, higher lipid oxidation in M(IL-4), seen with only GlyGln supplemented medium, is consistent with murine M(IL-4) metabolism, in which fatty acid oxidation is indispensable for IL-4-induced polarization (60).

In an earlier paper, we reported that the transcriptomic signature of human polarized MΦ was characterized by up-regulation of TCA cycle, ETC and oxidative phosphorylation genes in M(IL-4) and a mirror down-regulation in M(LPS+INF-γ) (9). In comparison to transcriptomic differences, the changes herein reported between M(LPS+INF-γ) and M(IL-4) are apparently minor. Altogether, this is an example of partial discrepancy between transcriptomic data and fluxes of central carbon metabolism under basal conditions and reinforces the rationale for measuring flux rates in the quest of MΦ metabolic signatures, as suggested by the principles of fluxomics (11).

Our study is not devoid of limitations.

First, ammonia quantification is critical to estimate amino acid catabolism. Human MΦ secrete a very low quantity of NH_4_, requiring a highly sensitive estimation method. We believe that there is significant room for improvement. Secondly, the estimates of amino acid catabolism rely on the composition of the “proteomoid” and on the relative contributions of the “proteomoid” and GlyGln to amino acid catabolism: both are only educated assumptions. Third, the total ATP production remains unknown, since we can compute only an upper boundary (maximum ATP production) and a lower boundary (ATP production due to anaerobic glycolysis). Fourth, this cell indirect microcalorimetry technique shares the same theoretical strengths and limitations of indirect calorimetry (17, 42). Fifth, monocyte-derived macrophage *in vitro* polarization represents an extremization and oversimplification of the *in vivo* micro-environment, characterized by a plethora of phenotypes (36). Sixth, in our experience there is a high donor-to-donor variability, which we tried to mitigate by increasing the number of replicates.

In summary, we have set up a novel microcalorimetric technique to investigate cell energetic fluxomics. In our hands, the energetic fluxomic signature of human primary MΦ was characterized by:

i. Abundant usage of glycogen/glucose to support a huge glycolytic flux and about 30% of the citrate synthase reaction, a rate limiting step being the PDH reaction
ii. Predominance of lipids in sustaining net citrate synthesis, i.e. the first step of TCA cycle Further patterns emerged when polarized MΦ were scrutinized:
iii. M(LPS+INF-γ) feature higher anaerobic glycolysis than M0 and M(IL-4), similar TCA cycle fluxes as M0
iv. With glucose and GlyGln supply, M(IL-4) unveil higher TCA cycle enzyme fluxes than M0, and higher malic enzyme flux than M1, plus a general trend to higher lipid oxidation rate with GlyGln supplemented media.

These data were obtained in human primary MΦ and show significant differences with the murine MΦ metabolism, but also similarities, e.g. higher anaerobic glycolytic flux in M(LPS+INF-γ), which thus far have been difficult to document.

We believe that, especially if combined with tracer techniques and/or quantitative metabolomics, this cell indirect microcalorimetry technique can unveil novel details of metabolic regulation in immune cells, and also in other cell systems.

## Acknowledgements

We thank Rosanna Vescovini at the University of Parma, Italy, for expert technical assistance in flow citometry.

This work was supported by the Italian Ministry of University and Research (MUR) through the PRIN 2020 grant n. 2020SH2ZZA_003 to RCB, and by the University of Parma through “Bando straordinario di Ateneo per progetti di ricerca biomedica in ambito SARS-CoV-2 e COVID-19 2020” to RCB and through “FIL” funds for research to ADC and RCB.

## Supplemental Information

Document S1. Supplementary Table 1-4, Figure S1 and S2, Supplemental Methods (Equation entered into the SAAM II Software)

## Author Contributions

*Conceived and designed research,* RCB; *Performed experiments,* GC, FF, ADA, FB and EG; *Analyzed data,* GC, AB and VS; *Interpreted results of experiments,* GC, VS, ADC, RCB; *Prepared figures,* GC; *Drafted manuscript,* GC and VS.; *Edited and revised the manuscript,* ADA, ADC and RCB; *Approved final version of the manuscript,* GC, VS, FF, ADA, FB, EG, AB, ADC and RCB.

