## Supplementary material for "A novel cell indirect calorimetry method unveils the metabolic fluxomic signatures of human monocyte derived M(LPS+INF-γ) and M(IL-4) macrophages"

**Supplementary Table 1. Equations of amino acid oxidation to pyruvate and CO_2_**

| **Amino acid** | **Oxidation to CO_2_** |
| --- | --- |
| Glutamine | $\boldsymbol{Gln+2}\boldsymbol{O}_{\boldsymbol{2}}\boldsymbol{+2}\boldsymbol{H}_{\boldsymbol{2}}\boldsymbol{O \to Py}\boldsymbol{r}^{\boldsymbol{-}}\boldsymbol{+2}\boldsymbol{N}\boldsymbol{H}_{\boldsymbol{4}}^{\boldsymbol{+}}\boldsymbol{+}\boldsymbol{H}^{\boldsymbol{+}}\boldsymbol{+2 HC}\boldsymbol{O}_{\boldsymbol{3}}^{\boldsymbol{-}}$  $\dot{\boldsymbol{OCR}}\boldsymbol{=}\dot{\boldsymbol{N}}\boldsymbol{+ 1.25\cdot}\left( \boldsymbol{1-x} \right)\boldsymbol{\cdot}\dot{\boldsymbol{N}}\boldsymbol{-0.25 x\cdot}\dot{\boldsymbol{N}}$  $\dot{\boldsymbol{PPR}}\boldsymbol{=0.5\cdot}\dot{\boldsymbol{N}}\boldsymbol{+}\left( \boldsymbol{1-x} \right)\boldsymbol{\cdot}\dot{\boldsymbol{N}}\boldsymbol{-0.5 x\cdot}\dot{\boldsymbol{N}}$ |
| Aspartate | $\boldsymbol{As}\boldsymbol{p}^{-}+\mathbf{0}.\mathbf{5} \boldsymbol{O}_{\mathbf{2}}+\boldsymbol{H}_{\mathbf{2}}\boldsymbol{O} \to\boldsymbol{Py}\boldsymbol{r}^{-}+\boldsymbol{N}\boldsymbol{H}_{\mathbf{4}}^{+}+ \boldsymbol{HC}\boldsymbol{O}_{\mathbf{3}}^{-}$  $\dot{\boldsymbol{OCR}}\boldsymbol{=0.5\cdot}\dot{\boldsymbol{N}}\boldsymbol{+2.5\cdot}\left( \boldsymbol{1-x} \right)\boldsymbol{\cdot}\dot{\boldsymbol{N}}\boldsymbol{-0.5 x\cdot}\dot{\boldsymbol{N}}$  $\dot{\boldsymbol{PPR}}\boldsymbol{=2\cdot}\left( \boldsymbol{1-x} \right)\boldsymbol{\cdot}\dot{\boldsymbol{N}}\boldsymbol{- x\cdot}\dot{\boldsymbol{N}}$ |
| Asparagine | $\boldsymbol{Asn}+\mathbf{0}.\mathbf{5} \boldsymbol{O}_{\mathbf{2}}+\mathbf{2}\boldsymbol{H}_{\mathbf{2}}\boldsymbol{O} \to\boldsymbol{Py}\boldsymbol{r}^{-}+\mathbf{2}\boldsymbol{N}\boldsymbol{H}_{\mathbf{4}}^{+}+ \boldsymbol{HC}\boldsymbol{O}_{\mathbf{3}}^{-}$  $\dot{\boldsymbol{OCR}}\boldsymbol{=0.25\cdot}\dot{\boldsymbol{N}}\boldsymbol{+1.25\cdot}\left( \boldsymbol{1-x} \right)\boldsymbol{\cdot}\dot{\boldsymbol{N}}\boldsymbol{-0.25 x\cdot}\dot{\boldsymbol{N}}$  $\dot{\boldsymbol{PPR}}\boldsymbol{=}\left( \boldsymbol{1-x} \right)\boldsymbol{\cdot}\dot{\boldsymbol{N}}\boldsymbol{- 0.5 x\cdot}\dot{\boldsymbol{N}}$ |
| Valine | $\boldsymbol{Val}+\mathbf{3}.\mathbf{5} \boldsymbol{O}_{\mathbf{2}} \to\boldsymbol{Py}\boldsymbol{r}^{-}+\boldsymbol{N}\boldsymbol{H}_{\mathbf{4}}^{+}+\mathbf{2} \boldsymbol{HC}\boldsymbol{O}_{\mathbf{3}}^{-}+ \mathbf{2}\boldsymbol{H}^{+}$  $\dot{\boldsymbol{OCR}}\boldsymbol{=3.5\cdot}\dot{\boldsymbol{N}}\boldsymbol{+2.5\cdot}\left( \boldsymbol{1-x} \right)\boldsymbol{\cdot}\dot{\boldsymbol{N}}\boldsymbol{-0.5 x\cdot}\dot{\boldsymbol{N}}$  $\dot{\boldsymbol{PPR}}\boldsymbol{=2}\dot{\boldsymbol{N}}\boldsymbol{+2}\left( \boldsymbol{1-x} \right)\boldsymbol{\cdot}\dot{\boldsymbol{N}}\boldsymbol{- x\cdot}\dot{\boldsymbol{N}}$ |
| Isoleucine | $\boldsymbol{Ile}+\mathbf{5} \boldsymbol{O}_{\mathbf{2}} \to\boldsymbol{Py}\boldsymbol{r}^{-}+\boldsymbol{N}\boldsymbol{H}_{\mathbf{4}}^{+}+\mathbf{3} \boldsymbol{HC}\boldsymbol{O}_{\mathbf{3}}^{-}+ \mathbf{3}\boldsymbol{H}^{+}$  $\dot{\boldsymbol{OCR}}\boldsymbol{=5\cdot}\dot{\boldsymbol{N}}\boldsymbol{+2.5\cdot}\left( \boldsymbol{1-x} \right)\boldsymbol{\cdot}\dot{\boldsymbol{N}}\boldsymbol{-0.5 x\cdot}\dot{\boldsymbol{N}}$  $\dot{\boldsymbol{PPR}}\boldsymbol{=3\cdot}\dot{\boldsymbol{N}}\boldsymbol{+2}\left( \boldsymbol{1-x} \right)\boldsymbol{\cdot}\dot{\boldsymbol{N}}\boldsymbol{- x\cdot}\dot{\boldsymbol{N}}$ |
| Alanine | $\boldsymbol{Ala}+{\mathbf{0}.\mathbf{5} \boldsymbol{O}}_{\mathbf{2}} \to\boldsymbol{Py}\boldsymbol{r}^{-}+\boldsymbol{N}\boldsymbol{H}_{\mathbf{4}}^{+}$  $\dot{\boldsymbol{OCR}}\boldsymbol{=0.5\cdot}\dot{\boldsymbol{N}}\boldsymbol{+ 2.5\cdot}\left( \boldsymbol{1-x} \right)\boldsymbol{\cdot}\dot{\boldsymbol{N}}\boldsymbol{-0.5 x\cdot}\dot{\boldsymbol{N}}$  $\dot{\boldsymbol{PPR}}\boldsymbol{=2}\left( \boldsymbol{1-x} \right)\boldsymbol{\cdot}\dot{\boldsymbol{N}}\boldsymbol{-x\cdot}\dot{\boldsymbol{N}}$ |
| Arginine | $\boldsymbol{Ar}\boldsymbol{g}^{+}+\mathbf{3}\boldsymbol{O}_{\mathbf{2}}+\mathbf{2}\boldsymbol{H}_{\mathbf{2}}\boldsymbol{O} \to\boldsymbol{Py}\boldsymbol{r}^{-}+\mathbf{2}\boldsymbol{N}\boldsymbol{H}_{\mathbf{4}}^{+}+ {\mathbf{2}\boldsymbol{H}}^{+}+ \mathbf{2}\boldsymbol{HC}\boldsymbol{O}_{\mathbf{3}}^{-}+\boldsymbol{Urea}$  $\dot{\boldsymbol{OCR}}\boldsymbol{=3}\dot{\boldsymbol{N}}\boldsymbol{+ 2.5\cdot}\left( \boldsymbol{1-x} \right)\boldsymbol{\cdot}\dot{\boldsymbol{N}}\boldsymbol{-0.5 x\cdot}\dot{\boldsymbol{N}}$  $\dot{\boldsymbol{PPR}}\boldsymbol{=2}\dot{\boldsymbol{N}}\boldsymbol{+2}\left( \boldsymbol{1-x} \right)\boldsymbol{\cdot}\dot{\boldsymbol{N}}\boldsymbol{-x\cdot}\dot{\boldsymbol{N}}$ |
| Cysteine | $\boldsymbol{Cys}+\mathbf{2}\boldsymbol{O}_{\mathbf{2}}+ \boldsymbol{H}_{\mathbf{2}}\boldsymbol{O} \to\boldsymbol{Py}\boldsymbol{r}^{-}+{\boldsymbol{SO}\mathbf{4}}^{\mathbf{2}-}+\boldsymbol{N}\boldsymbol{H}_{\mathbf{4}}^{+}+ {\mathbf{2}\boldsymbol{H}}^{+}$  $\dot{\boldsymbol{OCR}}\boldsymbol{=2}\dot{\boldsymbol{N}}\boldsymbol{+ 2.5\cdot}\left( \boldsymbol{1-x} \right)\boldsymbol{\cdot}\dot{\boldsymbol{N}}\boldsymbol{-0.5 x\cdot}\dot{\boldsymbol{N}}$  $\dot{\boldsymbol{PPR}}\boldsymbol{=2}\dot{\boldsymbol{N}}\boldsymbol{+2}\left( \boldsymbol{1-x} \right)\boldsymbol{\cdot}\dot{\boldsymbol{N}}\boldsymbol{-x\cdot}\dot{\boldsymbol{N}}$ |
| Glycine | $\mathbf{2}\boldsymbol{Gly}+\mathbf{0}.\mathbf{5}\boldsymbol{O}_{\mathbf{2}}+ \boldsymbol{H}_{\mathbf{2}}\boldsymbol{O}\to\boldsymbol{Py}\boldsymbol{r}^{-}+\mathbf{2}\boldsymbol{N}\boldsymbol{H}_{\mathbf{4}}^{+}+ \boldsymbol{HC}\boldsymbol{O}_{\mathbf{3}}^{-}$  $\dot{\boldsymbol{OCR}}\boldsymbol{=0.25\cdot N+ 1.25\cdot}\left( \boldsymbol{1-x} \right)\boldsymbol{\cdot}\dot{\boldsymbol{N}}\boldsymbol{-0.25 x\cdot}\dot{\boldsymbol{N}}$  $\dot{\boldsymbol{PPR}}\boldsymbol{=}\left( \boldsymbol{1-x} \right)\boldsymbol{\cdot}\dot{\boldsymbol{N -0.5\cdot x\cdot}\dot{\boldsymbol{N}}}$ |
| Glutamate | $\boldsymbol{Gl}\boldsymbol{u}^{-}+\mathbf{2}\boldsymbol{O}_{\mathbf{2}}+\boldsymbol{H}_{\mathbf{2}}\boldsymbol{O} \to\boldsymbol{Py}\boldsymbol{r}^{-}+\boldsymbol{N}\boldsymbol{H}_{\mathbf{4}}^{+}+ \boldsymbol{H}^{+}+\mathbf{2} \boldsymbol{HC}\boldsymbol{O}_{\mathbf{3}}^{-}$  $\dot{\boldsymbol{OCR}}\boldsymbol{=2}\dot{\boldsymbol{N}}\boldsymbol{+ 2.5\cdot}\left( \boldsymbol{1-x} \right)\boldsymbol{\cdot}\dot{\boldsymbol{N}}\boldsymbol{-0.5 x\cdot}\dot{\boldsymbol{N}}$  $\dot{\boldsymbol{PPR}}\boldsymbol{=}\dot{\boldsymbol{N}}\boldsymbol{+2}\left( \boldsymbol{1-x} \right)\boldsymbol{\cdot}\dot{\boldsymbol{N}}\boldsymbol{-x\cdot}\dot{\boldsymbol{N}}$ |
| Methionine | $\boldsymbol{Met}\mathbf{+3.5}\boldsymbol{O}_{\mathbf{2}}\mathbf{+}\boldsymbol{H}_{\mathbf{2}}\boldsymbol{O}\boldsymbol{\to}\boldsymbol{Py}\boldsymbol{r}^{\mathbf{-}}\mathbf{+}{\boldsymbol{SO}\mathbf{4}}^{\mathbf{2-}}\mathbf{+}\boldsymbol{N}\boldsymbol{H}_{\mathbf{4}}^{\mathbf{+}}\mathbf{+}{\mathbf{3}\boldsymbol{H}}^{\mathbf{+}}\mathbf{+}\boldsymbol{HC}\boldsymbol{O}_{\mathbf{3}}^{\mathbf{-}}\mathbf{+}\boldsymbol{CH}_{\mathbf{2}}$  $\dot{\boldsymbol{OCR}}\boldsymbol{=3.5}\dot{\boldsymbol{N}}\boldsymbol{+ 2.5\cdot}\left( \boldsymbol{1-x} \right)\boldsymbol{\cdot}\dot{\boldsymbol{N}}\boldsymbol{-0.5 x\cdot}\dot{\boldsymbol{N}}$  $\dot{\boldsymbol{PPR}}\boldsymbol{=3}\dot{\boldsymbol{N}}\boldsymbol{+2}\left( \boldsymbol{1-x} \right)\boldsymbol{\cdot}\dot{\boldsymbol{N}}\boldsymbol{-x\cdot}\dot{\boldsymbol{N}}$ |
| Serine | $\boldsymbol{Ser}\to\boldsymbol{Py}\boldsymbol{r}^{-}+\boldsymbol{N}\boldsymbol{H}_{\mathbf{4}}^{+}$  $\dot{\boldsymbol{OCR}}\boldsymbol{=2.5\cdot}\left( \boldsymbol{1-x} \right)\boldsymbol{\cdot}\dot{\boldsymbol{N}}\boldsymbol{-0.5 x\cdot}\dot{\boldsymbol{N}}$  $\dot{\boldsymbol{PPR}}\boldsymbol{=2}\left( \boldsymbol{1-x} \right)\boldsymbol{\cdot}\dot{\boldsymbol{N}}\boldsymbol{-x\cdot}\dot{\boldsymbol{N}}$ |
| Threonine | $\boldsymbol{Thr}+\mathbf{1}.\mathbf{5}\boldsymbol{O}_{\mathbf{2}}\to\boldsymbol{Py}\boldsymbol{r}^{-}+\boldsymbol{N}\boldsymbol{H}_{\mathbf{4}}^{+}+ \boldsymbol{H}^{+}+ \boldsymbol{HC}\boldsymbol{O}_{\mathbf{3}}^{-}$  $\dot{\boldsymbol{OCR}}\boldsymbol{=1.5}\dot{\boldsymbol{N}}\boldsymbol{+ 2.5\cdot}\left( \boldsymbol{1-x} \right)\boldsymbol{\cdot}\dot{\boldsymbol{N}}\boldsymbol{-0.5 x\cdot}\dot{\boldsymbol{N}}$  $\dot{\boldsymbol{PPR}}\boldsymbol{=}\dot{\boldsymbol{N}}\boldsymbol{+2}\left( \boldsymbol{1-x} \right)\boldsymbol{\cdot}\dot{\boldsymbol{N}}\boldsymbol{-x\cdot}\dot{\boldsymbol{N}}$ |
| Leucine | $\boldsymbol{Leu}+\mathbf{7}.\mathbf{5} \boldsymbol{O}_{\mathbf{2}} + \boldsymbol{H}_{\mathbf{2}}\boldsymbol{O} \to\boldsymbol{N}\boldsymbol{H}_{\mathbf{4}}^{+}+\mathbf{6} \boldsymbol{HC}\boldsymbol{O}_{\mathbf{3}}^{-}+ \mathbf{5}\boldsymbol{H}^{+}$  $\dot{\boldsymbol{OCR}}\boldsymbol{=7.5\cdot}\dot{\boldsymbol{N}}$  $\dot{\boldsymbol{PPR}}\boldsymbol{=5\cdot}\dot{\boldsymbol{N}}$ |
| Lysine | $\boldsymbol{Ly}\boldsymbol{s}^{+}+\mathbf{7} \boldsymbol{O}_{\mathbf{2}} +\mathbf{2} \boldsymbol{H}_{\mathbf{2}}\boldsymbol{O} \to\mathbf{2}\boldsymbol{N}\boldsymbol{H}_{\mathbf{4}}^{+}+\mathbf{6} \boldsymbol{HC}\boldsymbol{O}_{\mathbf{3}}^{-}+ \mathbf{5}\boldsymbol{H}^{+}$  $\dot{\boldsymbol{OCR}}\boldsymbol{=3.5\cdot}\dot{\boldsymbol{N}}$  $\dot{\boldsymbol{PPR}}\boldsymbol{=2.5\cdot}\dot{\boldsymbol{N}}$ |
| Tryptophan | $\boldsymbol{Trp}+\mathbf{9} \boldsymbol{O}_{\mathbf{2}} +\mathbf{4} \boldsymbol{H}_{\mathbf{2}}\boldsymbol{O} \to\boldsymbol{Py}\boldsymbol{r}^{-}+\mathbf{2}\boldsymbol{N}\boldsymbol{H}_{\mathbf{4}}^{+}+\mathbf{8} \boldsymbol{HC}\boldsymbol{O}_{\mathbf{3}}^{-}+ \mathbf{7}\boldsymbol{H}^{+}$  $\dot{\boldsymbol{OCR}}\boldsymbol{=4.5\cdot}\dot{\boldsymbol{N}}\boldsymbol{+ 1.25\cdot}\left( \boldsymbol{1-x} \right)\boldsymbol{\cdot}\dot{\boldsymbol{N}}\boldsymbol{-0.25 x\cdot}\dot{\boldsymbol{N}}$  $\dot{\boldsymbol{PPR}}\boldsymbol{=3.5\cdot}\dot{\boldsymbol{N}}\boldsymbol{+}\left( \boldsymbol{1-x} \right)\boldsymbol{\cdot}\dot{\boldsymbol{N}}\boldsymbol{-0.5 x\cdot}\dot{\boldsymbol{N}}$ |
| GlyGln | $\boldsymbol{GlyGln+2.25}\boldsymbol{O}_{\boldsymbol{2}}\boldsymbol{+3.5}\boldsymbol{H}_{\boldsymbol{2}}\boldsymbol{O \to1.5}\boldsymbol{Py}\boldsymbol{r}^{\boldsymbol{-}}\boldsymbol{+3}\boldsymbol{N}\boldsymbol{H}_{\boldsymbol{4}}^{\boldsymbol{+}}\boldsymbol{+}\boldsymbol{H}^{\boldsymbol{+}}\boldsymbol{+2.5 HC}\boldsymbol{O}_{\boldsymbol{3}}^{\boldsymbol{-}}$  $\dot{\boldsymbol{OCR}}\boldsymbol{=}\dot{\boldsymbol{1.125}\boldsymbol{N}}\boldsymbol{+ 1.875\cdot}\left( \boldsymbol{1-x} \right)\boldsymbol{\cdot}\dot{\boldsymbol{N}}\boldsymbol{-0.375 x\cdot}\dot{\boldsymbol{N}}$  $\dot{\boldsymbol{PPR}}\boldsymbol{=0.5\cdot}\dot{\boldsymbol{N}}\boldsymbol{+1.5\cdot}\left( \boldsymbol{1-x} \right)\boldsymbol{\cdot}\dot{\boldsymbol{N}}\boldsymbol{-0.75 x\cdot}\dot{\boldsymbol{N}}$ |

Equations of complete amino acid oxidation and their contribution to oxygen consumption ($O\dot{C}R$) and proton production ($P\dot{P}R$) expressed as a function of ammonia fluxes $(\dot{N})$. $O\dot{C}R$= molecular oxygen consumption rate; $P\dot{P}R$ = proton production rate; $\dot{N}$ = NH_4_ flux; Gly= glycine; Ala= alanine; Val= valine; Leu= leucine; Ile = isoleucine; Met= methionine; Cys= cysteine; Trp= tryptophan; Lys= lysine; Arg= arginine; Gln= glutamine; Asn= asparagine; Glu= glutamate; Asp= aspartate; Ser= serine; Thr= threonine; GlyGln= Glycyl-glutamine dipeptide.

**Supplementary Table 2. ATP production due to amino acid catabolism.**

| **Amino acid** | **ATP flux** |
| --- | --- |
| Glutamine | $A\dot{T}P=5.28\cdot\dot{N}+6.65 \left( 1-x \right)\cdot\dot{N}- 1.36 x\cdot\dot{N}$ |
| Aspartate | $A\dot{T}P=2.73\cdot\dot{N}+13.29 \left( 1-x \right)\cdot\dot{N}- 2.73 x\cdot\dot{N}$ |
| Asparagine | $A\dot{T}P=1.36\cdot\dot{N}+6.65 \left( 1-x \right)\cdot\dot{N}- 1.36 x\cdot\dot{N}$ |
| Valine | $A\dot{T}P=16.93\cdot\dot{N}+13.29 \left( 1-x \right)\cdot\dot{N}- 2.73 x\cdot\dot{N}$ |
| Isoleucine | $A\dot{T}P=24.76\cdot\dot{N}+13.29 \left( 1-x \right)\cdot\dot{N}- 2.73 x\cdot\dot{N}$ |
| Alanine | $A\dot{T}P= 2.73\cdot\dot{N}+13.29 \left( 1-x \right)\cdot\dot{N}- 2.73 x\cdot\dot{N}$ |
| Arginine | $A\dot{T}P= 8.01\cdot\dot{N}+6.65 \left( 1-x \right)\cdot\dot{N}- 1.36 x\cdot\dot{N}$ |
| Cysteine | $A\dot{T}P= 2.73\cdot\dot{N}+13.29 \left( 1-x \right)\cdot\dot{N}- 2.73 x\cdot\dot{N}$ |
| Glycine | $A\dot{T}P= 1.36\cdot\dot{N}+6.65 \left( 1-x \right)\cdot\dot{N}- 1.36 x\cdot\dot{N}$ |
| Glutamate | $A\dot{T}P= 10.56\cdot\dot{N}+13.29 \left( 1-x \right)\cdot\dot{N}- 2.73 x\cdot\dot{N}$ |
| Methionine | $A\dot{T}P= 8.83\cdot\dot{N}+13.29 \left( 1-x \right)\cdot\dot{N}- 2.73 x\cdot\dot{N}$ |
| Serine | $A\dot{T}P= 13.29 \left( 1-x \right)\cdot\dot{N}- 2.73 x\cdot\dot{N}$ |
| Threonine | $A\dot{T}P= 7.1\cdot\dot{N}+13.29 \left( 1-x \right)\cdot\dot{N}- 2.73 x\cdot\dot{N}$ |
| Leucine | $A\dot{T}P= 37.32\cdot\dot{N}$ |
| Lysine | $A\dot{T}P= 17.84\cdot\dot{N}$ |
| Tryptophan | $A\dot{T}P= 13.38\cdot\dot{N}+6.65 \left( 1-x \right)\cdot\dot{N}- 1.36 x\cdot\dot{N}$ |
| GlyGln | $A\dot{T}P=6.64\cdot\dot{N}+9.975 \left( 1-x \right)\cdot\dot{N}- 2.04 x\cdot\dot{N}$ |

The table reports the equations relative to amino acid contribution to ATP flux $(A\dot{T}P$) as equivalent of ammonia flux ($\dot{N}$). $\dot{N}$ = NH_4_ flux;

**Supplemental Methods**

**Equations entered into the SAAM II software to calculate fluxomic parameters.**

The measured (known) parameters are lactate release rate (LDH), oxygen consumption (OCR), proton production rate (PPR) and NH_4_ release rate (N). The unknown parameters estimated by SAAM 2 are *ṁG* (irreversible glucose utilization rate), *ṁL* (net lipid oxidation rate, *PDH* (total flux of the PDH reaction), and *ω* (the fraction of protons generated by anaerobic glycolysis which is not buffered by other reactions, e.g. adenylate kinase).

**SUBSTRATE DEPRIVED MEDIUM**

*x* = LDH /(LDH + *PDH*)

OCR= 6*(1*-x*)**ṁG* + 75**ṁL* + 3.5*(1-*x*)**ṁL* + 2.247*N + 1.56*(1-*x*)*N - 0.312**x**N

PPR= 6*(1-*x*)**ṁG* + *ω***x***ṁG* + 52**ṁL* + 2*(1-*x*)**ṁL* - *x**ṁL + 1.426*N + 1.248*(1-*x*)*N - 0.596**x**N

NH_4_ Input into Acetyl-CoA= 0.451*N

NH_4_ Input into αKG= 0.167*N

NH_4_ Input into Succinyl-CoA= 0.1698*N

NH_4_ Input into Pyr= 0.208*N

Acetyl-CoA Input= *PDH* + 26*ṁL + NH_4_ Input into Acetyl-CoA

Net Citrate Synthase Flux= 26**ṁL* + *PDH* + NH_4_ Input into Acetyl-CoA

Aconitase= 26**ṁL* + *PDH* + NH_4_ Input into Acetyl-CoA

Net TCA Flux= Aconitase + 0.5*NH_4_ Input into αKG

αKGDH= Aconitase + NH_4_ Input into αKG

SDH= αKGDH + NH_4_ Input into SA

Malate Turnover= SDH + 0.0791*N

Malic Enzyme= Malate Turnover - Aconitase

Malate minus Oxaloacetate turnover= Malate Turnover – Net Citrate Synthase Flux

Pyruvate Turnover= 2**ṁG* + *ṁL* + NH_4_ Input into Pyr + MalicEnzyme

Anaerobic Glycolysis= [(2**ṁG*/Pyruvate Turnover)*LDH]/2

Glucose Oxidation= ṁG – Anaerobic Glycolysis

EPR= -2.733*(1-x)*ṁG - 0.115*x*ṁG - 33.25*ṁL + 1.1*x*ṁL - 0.9072*N - 0.7094*(1-x)*N - 0.04825*x*N

Anaerobic EPR= -0.115*x*ṁG - 0.5536*x*ṁL - 0.04825*x*N

Max ATP production= 33.5*(1-x)*ṁG + 2.9*x*ṁG + 367*ṁL + 19.3*(1-x)*ṁL + x*ṁL + 10.807*N + 8.295*(1-x)*N - 1.702*x*N

P/O= Max ATP production/(2*OCR)

Energy Efficiency= -(0.03155* Max ATP production)/(EPR-1.3088*LDH)

Anaerobic ATP= 2.9*x*ṁG + x*ṁL

Aerobic P/O= (Max ATP production – Anaerobic ATP)/(2*OCR-0.101*N)

Aerobic Energy Efficiency= -[0.03155*(Max ATP production –Anaerobic ATP)]/(EPR – Anaerobic EPR)

**GLUCOSE (5 mM) MEDIUM**

*x* = LDH /(LDH + *PDH*)

OCR= 6*(1-x)*ṁG + 75*ṁL + 3.5*(1-x)*ṁL + 2.247*N + 1.56*(1-x)*N - 0.312*x*N

PPR= 6*(1-x)*ṁG + *ω**x*ṁG + 52*ṁL + 2*(1-x)*ṁL - x*ṁL + 1.426*N + 1.248*(1-x)*N - 0.596*x*N

NH_4_ Input into Acetyl-CoA= 0.451*N

NH_4_ Input into αKG= 0.167*N

NH_4_ Input into Succinyl-CoA= 0.1698*N

NH_4_ Input into Pyr= 0.208*N

Acetyl-CoA Input= PDH + 26*ṁL + NH_4_ Input into Acetyl-CoA

Net Citrate Synthase flux= 26*ṁL + PDH + NH_4_ Input into Acetyl-CoA

Aconitase= 26*ṁL + PDH + NH_4_ Input into Acetyl-CoA

Net TCA Flux =Aconitase + 0.5* NH_4_ Input into αKG

αKGDH= Aconitase + NH_4_ Input into αKG

SDH= αKGDH + NH_4_ Input into SA

Malate Turnover= SDH + 0.0791*N

Malic Enzyme= Malate Turnover - Aconitase

Malate minus Oxaloacetate turnover= Malate Turnover – Net Citrate Synthase flux

Pyruvate turnover= 2*ṁG + ṁL + NH_4_ Input into Pyr + Malic Enzyme

Anaerobic Glycolysis= [(2*ṁG/Pyruvate turnover)*LDH]/2

Glucose Oxidation= ṁG – Anaerobic Glycolysis

EPR= -2.733*(1-x)*ṁG - 0.115*x*ṁG - 33.25*ṁL + 1.31*x*ṁL - 0.9072*N - 0.7094*(1-x)*N - 0.04825*x*N

Anaerobic EPR= -0.115*x*ṁG - 0.5536*x*ṁL - 0.04825*x*N

Max ATP production= 32.65*(1-x)*ṁG + 2.0*x*ṁG + 367*ṁL + 19.3*(1-x)*ṁL + x*ṁL + 10.807*N + 8.295*(1-x)*N - 1.702*x*N

P/O= Max ATP production/(2*OCR)

Energy Efficiency= -(0.03155*Max ATP Production)/(EPR - 1.3088*LDH)

Anaerobic ATP= 2.0*x*ṁG + x*ṁL

Aerobic P/O= (Max ATP production – Anaerobic ATP)/(2*OCR - 0.101*N)

Aerobic Energy Efficiency= -[0.03155 * (Max ATP production –Anaerobic ATP)]/(EPR – Anaerobic EPR)

**GLYCYL-GLUTAMINE (2 mM) MEDIUM**

*x* = LDH /(LDH + *PDH*)

OCR= 6*(1-x)*ṁG + 75*ṁL + 3.5*(1-x)*ṁL + (1/4)*(2.247*N + 1.56*(1-x)*N - 0.312*x*N) + (3/4)*[0.75*N + 1.25*(1-x)*N - 0.25*x*N]

PPR= 6*(1-x)*ṁG + *ω**x*ṁG + 52*ṁL + 2*(1-x)*ṁL - x*ṁL + (1/4)*[1.426*N + 1.248*(1-x)*N - 0.596*x*N] + (3/4)*[0.333*N + (1-x)*N - 0.5*N]

NH_4_ Input into Acetyl-CoA= (1/4)*(0.451*N)

NH_4_ Input into αKG= (1/4)*(0.167*N) + (3/4)*(0.333*N)

NH_4_ Input into Succinyl-CoA= (1/4)*(0.1698*N)

NH_4_ Input into Pyr= (1/4)*(0.208*N) + (3/4)*(0.1667*N)

Acetyl-CoA Input= PDH + 26*ṁL + NH_4_ Input into Acetyl-CoA

Net Citrate Synthase flux= 26*ṁL + PDH + NH_4_ Input into Acetyl-CoA

Aconitase= 26*ṁL + PDH + NH_4_ Input into Acetyl-CoA

Net TCA Flux= Aconitase + 0.5* NH_4_ Input into αKG

αKGDH= Aconitase + NH_4_ Input into αKG

SDH= αKGDH+ NH_4_ Input into SA

Malate Turnover= SDH + 0.0791*N

Malic Enzyme= Malate Turnover - Aconitase

Malate minus Oxaloacetate turnover= Malate Turnover – Net Citrate Synthase flux

Pyruvate turnover= 2*ṁG + ṁL + NH_4_ Input into Pyr + MalicEnzyme

Anaerobic Glycolysis= ((2*ṁG/Pyruvate turnover)*LDH)/2

Glucose Oxidation= ṁG – Anaerobic Glycolysis

EPR= -2.733*(1-x)*ṁG - 0.115*x*ṁG - 33.25*ṁL + 1.1*x*ṁL - (1/4)*[0.9072*N + 0.7094*(1-x)*N + 0.04825*x*N] - (3/4)*[0.204*N + 0.568*(1-x)*N + 0.0387*x*N]

Anaerobic EPR= -0.115*x*ṁG - 0.5536*x*ṁL - (1/4)*0.04825*x*N - (3/4)*0.0387*x*N

Max ATP production= 33.5*(1-x)*ṁG + 2.9*x*ṁG + 367*ṁL + 19.3*(1-x)*ṁL + x*ṁL + (1/4)*[10.807*N + 8.295*(1-x)*N - 1.702*x*N] + (3/4)*[3.973*N + 6.65*(1-x)*N - 1.36*x*N]

P/O= Max ATP production/(2*OCR)

Energy Efficiency= -(0.03155* Max ATP production)/(EPR - 1.3088*LDH)

Anaerobic ATP= 2.9*x*ṁG + x*ṁL

Aerobic P/O= (Max ATP production – Anaerobic ATP)/(2*OCR - 0.101*N)

Aerobic Energy Efficiency= -[0.03155*(Max ATP production – Anaerobic ATP)]/(EPR – Anaerobic EPR)

**GLUCOSE (5 mM) + GLYCYL-GLUTAMINE (2 mM) MEDIUM**

*x* = LDH /(LDH + *PDH*)

OCR= 6*(1-x)*ṁG + 75*ṁL + 3.5*(1-x)*ṁL + (1/4)*[2.247*N + 1.56*(1-x)*N - 0.312*x*N] + (3/4)*[0.75*N + 1.25*(1-x)*N - 0.25*x*N]

PPR= 6*(1-x)*ṁG + *ω**x*ṁG + 52*ṁL + 2*(1-x)*ṁL - x*ṁL + (1/4)*[1.426*N + 1.248*(1-x)*N - 0.596*x*N] + (3/4)*[0.333*N + (1-x)*N - 0.5*N]

NH_4_ Input into Acetyl-CoA= (1/4)*(0.451*N)

NH_4_ Input into αKG= (1/4)*(0.167*N) + (3/4)*(0.333*N)

NH_4_ Input into Succinyl-CoA= (1/4)*(0.1698*N)

NH_4_ Input into Pyr= (1/4)*(0.208*N)+(3/4)*(0.1667*N)

Acetyl-CoA Input= PDH + 26*ṁL + NH_4_ Input into Acetyl-CoA

Net Citrate Synthase flux= 26*ṁL + PDH + NH_4_ Input into Acetyl-CoA

Aconitase= 26*ṁL + PDH + NH_4_ Input into Acetyl-CoA

Net TCA Flux= Aconitase + 0.5* NH_4_ Input into αKG

αKGDH= Aconitase + NH_4_ Input into αKG

SDH= αKGDH + NH_4_ Input into SA

Malate Turnover= SDH + 0.0791*N

Malic Enzyme= Malate Turnover - Aconitase

Malate minus Oxaloacetate turnover= Malate Turnover – Net Citrate Synthase flux

Pyruvate turnover= 2*ṁG + ṁL + NH_4_ Input into Pyr + Malic Enzyme

Anaerobic Glycolysis= [(2*ṁG/Pyruvate turnover)*LDH]/2

Glucose Oxidation= ṁG – Anaerobic Glycolysis

EPR= -2.733*(1-x)*ṁG - 0.115*x*ṁG - 33.25*ṁL + 1.1*x*ṁL - (1/4)*[0.9072*N + 0.7094*(1-x)*N + 0.04825*x*N] - (3/4)*[0.204*N + 0.568*(1-x)*N + 0.0387*x*N]

Anaerobic EPR= -0.115*x*ṁG - 0.5536*x*ṁL - (1/4)*0.04825*x*N - (3/4)*0.0387*x*N

Max ATP production= 32.65*(1-x)*ṁG + 2.0*x*ṁG + 367*ṁL + 19.3*(1-x)*ṁL + x*ṁL + (1/4)*[10.807*N + 8.295*(1-x)*N - 1.702*x*N] + (3/4)*[3.973*N + 6.65*(1-x)*N - 1.36*x*N]

P/O= Max ATP production/(2*OCR)

Energy Efficiency= -(0.03155*Max ATP production)/(EPR - 1.3088*LDH)

Anaerobic ATP= 2.0*x*ṁG + x*ṁL

Aerobic P/O= (Max ATP production – Anaerobic ATP)/(2*OCR-0.101*N)

Aerobic Energy Efficiency= -[0.03155*(Max ATP production – Anaerobic ATP)]/(EPR – Anaerobic EPR)

These data provide the equations used for calculation of fluxomic parameters related to substrate deprived medium, medium with glucose 5 mM, medium with glycyl-glutamine 2 mM and with both glucose + glycyl-glutamine. N= NH_4_ flux, LDH= lactate dehydrogenase flux, OCR= oxygen consumption flux, PPR= proton production flux, ṁG= glucose (or glycogen in glucose equivalents) utilization flux, ṁL= lipid oxidation flux, x= pyruvate contribution to lactate production, ω= the H+ buffering component due to intracellular (e.g. adenylate kinase) reactions, PDH= pyruvate dehydrogenase flux, αKG= α-ketoglutarate, αKGDH= α-ketoglutarate dehydrogenase flux, SDH= succinate dehydrogenase flux, EPR= energy production flux.

**Equations used in this study to calculate fluxomic parameters under conditions of net fatty acid synthesis (entered into the SAAM II software).**

The measured (known) parameters are lactate release rate (LDH), oxygen consumption rate (OCR), proton production rate (PPR) and NH_4_ release rate (N). The unknown parameters estimated by SAAM 2 are *ṁG* (irreversible glucose utilization rate), *Citrate Lyase* (flux of citrate lyase reaction), *PDH* (flux of the PDH reaction), and *ω* (the fraction of protons generated by anaerobic glycolysis, which is not buffered by other reactions, e.g. adenylate kinase).

**SUBSTRATE DEPRIVED MEDIUM**

x = LDH/(LDH + PDH)

ṁFS = Citrate Lyase/26

y = Citrate Lyase / (Citrate Lyase + Aconitase)

NH_4_ Input into Acetyl-CoA = 0.451*N

NH_4_ Input into αKG = 0.167*N

NH_4_ Input into Succinyl-CoA = 0.1698*N

NH_4_ Input into pyr = 0.208*N

Acetyl-CoA Input = PDH + NH_4_ Input into Acetyl-CoA

Net Citrate Synthase flux = PDH + NH_4_ Input into Acetyl-CoA

Aconitase = Net Citrate Synthase flux – Citrate Lyase

Net TCA Flux = Aconitase + 0.5* NH_4_ Input into αKG

αKGDH= Aconitase + NH_4_ Input into αKG

SDH= αKGDH + NH_4_ Input into Succinyl-CoA

Malate Turnover =SDH + 0.0791*N

Malic Enzyme = Malate Turnover - Aconitase

PPP = 6.7*ṁFS

Pyruvate Input = 2*(ṁG- PPP) + NH_4_ Input into pyr + Malic Enzyme + PPP - ṁFS

Malate minus Oxaloacetate turnover = Aconitase -Net Citrate Synthase flux

Anaerobic Glycolysis = (((2*ṁG - PPP - ṁFS)/ Pyruvate Input) * LDH)/2

Glucose Oxidation = (ṁG - Anaerobic Glycolysis) *(1-y) + (1/3)*y*(ṁG - Anaerobic Glycolysis)

Glucose Oxidation through PPP = PPP - 0.5*ṁFS - 0.5*x*(PPP - ṁFS)

Min G6PDH flux = 3*PPP

kFS from G = (((2*ṁG - PPP)/Pyruvate Input) *PDH)/Acetyl CoA Input

OCR = 2*(1-x)*(ṁG-PPP) + 4*(1-x)*(1-y)*(ṁG-PPP) + 0.5*(1-x)*PPP + 2*(1-x)*(1-y)*PPP + ṁFS + 1.276*N + 0.312*(1-x)*N + 1.248*(1-x)*(1-y)*N + 0.971*(1-y)*N-0.3387*x*N - 23*kFS from G*ṁFS - 20.23*(1-kFS from G)* ṁFS

Oxygen Utilization = 2*(1-x)*(ṁG-PPP) + 4*(1-x)*(1-y)*(ṁG-PPP) + 0.5*(1-x)*PPP + 2*(1-x)*(1-y)*PPP + 1.276*N + 0.312*(1-x)*N + 1.248*(1-x)*(1-y)*N + 0.971*(1-y)*N + ṁFS - 0.3387*x*N

PPR = 2*(1-x)*(ṁG-PPP) + 4*(1-x)*(1-y)*(ṁG-PPP) + *ω**x*(ṁG-PPP) + 1.072*N + 1.248*(1-x)*(1-y)*N + 0.395*(1-y)*N -0.596*x*N + 37*ṁFS

EPR = -0.19751*(1-x)*(ṁG-PPP) - 2.535*(1-x)*(1-y)*(ṁG-PPP) - 0.115*x*(ṁG-PPP) -0.2576*(1-x)*PPP - 0.9169*(1-x)*(1-y)*PPP -0.077*x*PPP - 0.434*N - 0.1607*(1-x)*N -0.572*(1-x)*(1-y)*N - 0.14*(1-y)*N -0.04825*x*N + 10.557*(1-kFS from G)*ṁFS + 8.211*kFS from G*ṁFS

Anaerobic EPR = -0.115*x*(ṁG-PPP) -0.077*x*PPP - 0.04825*x*N

Max ATP Production = 8.36*(1-x)*(ṁG-PPP) + 26.07*(1-x)*(1-y)*(ṁG-PPP) + 2.9*x*(ṁG-PPP) + 6.335*N + 1.702*(1-x)*N + 6.896*(1-x)*(1-y)*N + 4.451*(1-y)*N -1.702*x*N + 2.73*(1-x)*PPP + 10.54*(1-x)*(1-y)*PPP + 2*PPP

P/O = Max ATP Production/(2*OCR)

Energy Efficiency = -(0.03155*Max ATP Production)/(EPR - 1.3088*LDH)

Anaerobic ATP = 2.9*x*ṁG + 2*PPP

Aerobic P/O = (Max ATP Production -Anaerobic ATP)/(2*Oxygen Utilization - 0.101*N)

Aerobic Energy Efficiency = -(0.03155*(Max ATP Production – Anaerobic ATP))/(EPR-Anaerobic EPR)

**GLUCOSE (5 mM) MEDIUM**

x = LDH/(LDH + PDH)

ṁFS = Citrate Lyase/26

y = Citrate Lyase/(Citrate Lyase + Aconitase)

NH_4_ Input into Acetyl-CoA = 0.451*N

NH_4_ Input into αKG = 0.167*N

NH_4_ Input into Succinil-CoA = 0.1698*N

NH_4_ Input into Pyr = 0.208*N

Acetyl-CoA Input = PDH + NH_4_ Input into Acetyl-CoA

Net Citrate Synthase flux = PDH + NH_4_ Input into Acetyl-CoA

Aconitase = Net Citrate Synthase flux - Citrate Lyase

Net TCA Flux = Aconitase + 0.5* NH_4_ Input into αKG

αKGDH = Aconitase + NH_4_ Input into αKG

SDH = αKGDH + NH_4_ Input into Succinyl-CoA

Malate Turnover = SDH + 0.0791*N

Malic Enzyṁe = Malate Turnover - Aconitase

PPP = 6.7*ṁFS

Pyruvate Input = 2*(ṁG - PPP) + NH_4_ Input into Pyr + Ṁalic Enzyṁe + PPP -ṁFS

Malate minus Oxaloacetate turnover = Aconitase - Net Citrate Synthase flux

Anaerobic Glycolysis = (((2*ṁG – PPP - ṁFS)/Pyruvate Input)*LDH)/2

Glucose Oxidation = (ṁG – Anaerobic Glycolysis)*(1-y) + (1/3)*y*(ṁG – Anaerobic Glycolysis)

Glucose Oxidation through PPP = PPP - 0.5*ṁFS - 0.5*x*(PPP - ṁFS)

Min G6PDH flux = 3*PPP

kFS from G = (((2*ṁG -1*PPP)/Pyruvate Input)*PDH)/Acetyl-CoA Input

OCR = 2*(1-x)*(ṁG-PPP) + 4*(1-x)*(1-y)*(ṁG-PPP) + 0.5*(1-x)*PPP + 2*(1-x)*(1-y)*PPP + ṁFS + 1.276*N + 0.312*(1-x)*N + 1.248*(1-x)*(1-y)*N + 0.971*(1-y)*N - 0.3387*x*N - 23*kFS from G*ṁFS - 20.23*(1-kFS from G)*ṁFS

Oxygen Utilization = 2*(1-x)*(ṁG-PPP) + 4*(1-x)*(1-y)*(ṁG-PPP) + 0.5*(1-x)*PPP + 2*(1-x)*(1-y)*PPP + 1.276*N + 0.312*(1-x)*N + 1.248*(1-x)*(1-y)*N + 0.971*(1-y)*N + ṁFS - 0.3387*x*N

PPR=2*(1-x)*(ṁG-PPP) + 4*(1-x)*(1-y)*(ṁG-PPP) + *ω**x*(ṁG-PPP) + 1.072*N + 1.248*(1-x)*(1-y)*N + 0.395*(1-y)*N - 0.596*x*N + 37*ṁFS

EPR = -0.19751*(1-x)*(ṁG-PPP) - 2.535*(1-x)*(1-y)*(ṁG-PPP) - 0.115*x*(ṁG-PPP) - 0.2576*(1-x)*PPP - 0.9169*(1-x)*(1-y)*PPP - 0.077*x*PPP - 0.434*N - 0.1607*(1-x)*N - 0.572*(1-x)*(1-y)*N - 0.14*(1-y)*N - 0.04825*x*N + 10.557*(1-kFS from G)*ṁFS + 8.211*kFS from G*ṁFS

Anaerobic EPR = -0.115*x*(ṁG-PPP) - 0.077*x*PPP - 0.04825*x*N

Max ATP Production=7.46*(1-x)*(ṁG-PPP) + 25.17*(1-x)*(1-y)*(ṁG-PPP) + 2.0*x*(ṁG-PPP) + 6.335*N + 1.702*(1-x)*N + 6.896*(1-x)*(1-y)*N + 4.451*(1-y)*N - 1.702*x*N + 2.73*(1-x)*PPP + 10.54*(1-x)*(1-y)*PPP + PPP

P/O = Max ATP Production/(2*OCR)

Energy Efficiency = -(0.03155*Max ATP Production)/(EPR-1.3088*LDH)

Anaerobic ATP = 2.0*x*ṁG + 1*PPP

Aerobic P/O = (Max ATP Production – Anaerobic ATP)/(2*Oxygen Utilization - 0.101*N)

Aerobic Energy Efficiency = -(0.03155*(Max ATP Production – Anaerobic ATP))/(EPR-Anaerobic EPR)

These data provide the equations used for calculation of fluxomic parameters related to substrate deprived medium and medium with glucose 5 mM under conditions of net fatty acid synthesis. N= NH_4_ flux, LDH= lactate dehydrogenase flux, OCR= oxygen consumption flux, PPR= proton production flux, FS=fatty synthesis, ṁG= glucose (or glycogen in glucose equivalents) utilization flux, ṁL= lipid oxidation flux, x= pyruvate contribution to lactate production, ω= the H+ buffering component due to intracellular (e.g. adenylate kinase) reactions, PDH= pyruvate dehydrogenase flux, αKG= α-ketoglutarate, αKGDH= α-ketoglutarate dehydrogenase flux, SDH= succinate dehydrogenase flux, PPP= pentose phosphate pathway, G6PDH= glucose-6-phosphate dehydrogenase, EPR= energy production flux.

**Figure S1. Compartmental diagram of the metabolic model used to quantify the metabolic fluxes of human primary macrophages, as it is depicted in the SAAM compartmental modeling software (modified screenshot from the SAAM 2 software)**.

Lactate, oxygen, protons and NH4 compartments contain 0 mass at time=0 min and masses equal to their measured release rates per minute (lactate, protons, and NH4) or to their measured consumption rate per minute (oxygen) at time = 1 min. The syringe symbols represent fluxes into the compartments. Each of them is equal to the equations for OCR, PPR, Pyruvate Turnover, 26*mL and NH4 Input into Acetyl-CoA, as specified in the text of this Supplementary Information. The SAAM is used as a calculator, by letting it fit exactly the four measured fluxes (lactate, proton and NH4 release rates plus oxygen consumption rate) and thereby estimate the four unknown parameters (irreversible glucose/glycogen utilization rate, net lipid oxidation or synthesis rate, total flux of the PDH reaction and the fraction of protons generated by anaerobic glycolysis which is not buffered by other reactions) and all the other derived metabolic fluxes, under the assumption of metabolic steady state.

| Fat Synthesis  Parameters | Substrate deprived medium | | | Glucose | |
| --- | --- | --- | --- | --- | --- |
|  | **M0**  **N=1** | **M(IFN-γ+LPS)**  **N=2** | **M(IL-4)**  **N=1** | **M0**  **N=1** | **M(IFN-γ+LPS)**  **N=1** |
| Cytrate Lyase  (pmol/min) | 15.2 | 44.5±12.6 | 12.6 | 18.9 | 118 |
| Glucose Oxidation through Pentose Phosphate Pathway (pmol/min) | 2.2 | 6.6±2.0 | 1.7 | 2.5 | 16.5 |
| Pentose Phosphate Pathway (pmol/min) | 3.9 | 11.5±3.2 | 3.3 | 4.9 | 30.4 |
| Fat synthesis index  (pmol/min) | 0.6 | 1.7±0.5 | 0.5 | 0.7 | 4.5 |
| G6P-Dehydrogenase flux  (pmol/min) | 11.8 | 34.4±9.7 | 9.8 | 14.6 | 91.2 |

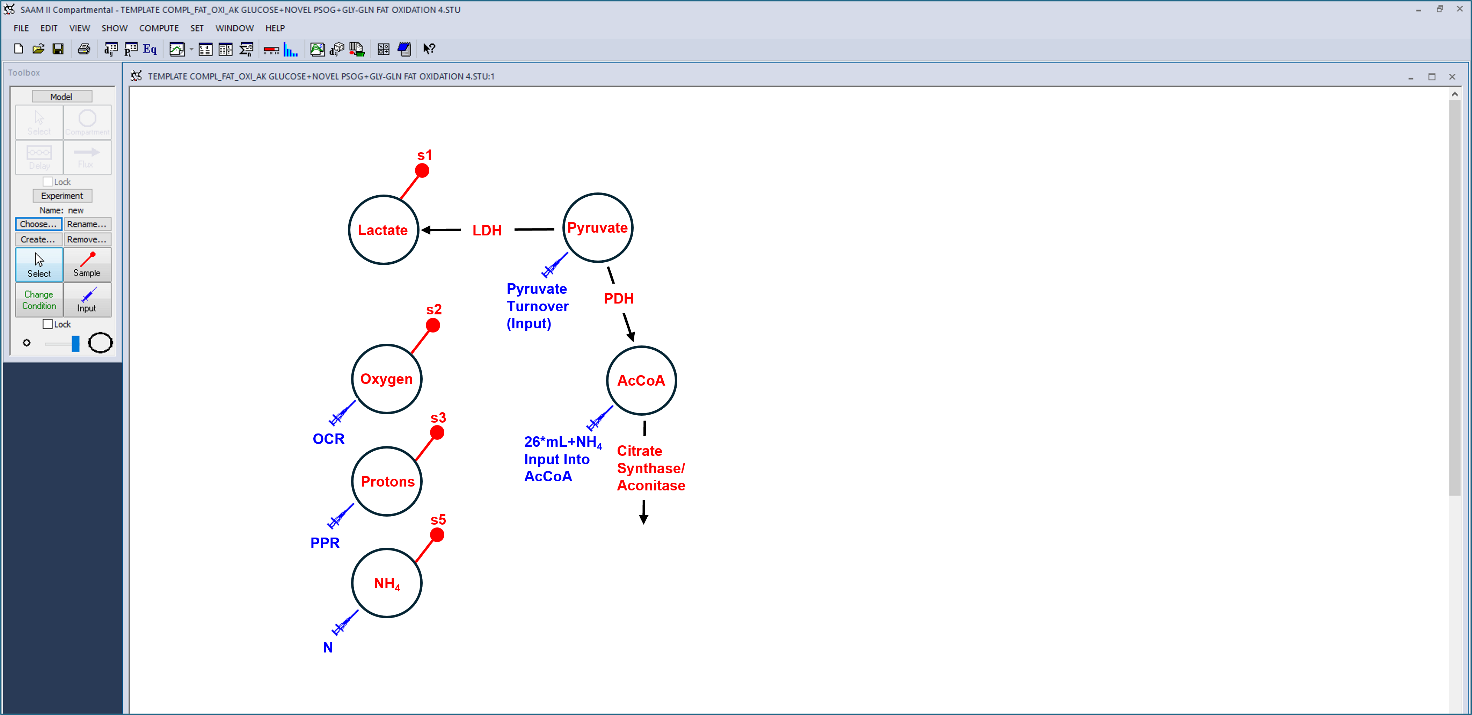

**Supplementary table 3. Fat Synthesis parameters in no exogenous substrates and glucose culture conditions.**

**Supplementary table 4. Mitostress test-derived respiratory parameters in M0, M(LPS+IFN-γ) and M(IL-4).**

|  | M0 | M(LPS+IFN-γ) | M(IL-4) |
| --- | --- | --- | --- |
| Basal respiration | 51.5±7.9 | 52.3±8.8 | 58.4±11.9 |
| ATP-linked respiration | 37.9±7.0 | 38.8±8.5 | 42.4±11.5 |
| Maximal respiration | 41.7±18.9 | 44.4±15.1 | 60.6±31.2 |
| Proton Leak | 13.6±3.7 | 13.5±1.1 | 16.0±2.7 |
| Spare respiratory capacity % | 74.3±29.1 | 79.5±22.9 | 87.1±29.5 |
| non-mitochondrial respiration | 18.0±1.6 | 18.8±1.3 | 19.3±2.7 |

The table reports four independent mitostress tests in which M0, M(LPS+INF-γ) and M(IL-4) macrophages derived from the same donor were tested in parallel. All parameters are reported as mean±SEM.

**Figure S2. Metabolic map used when lipid oxidation rate is < 0, i.e. indicative of net lipid synthesis.**

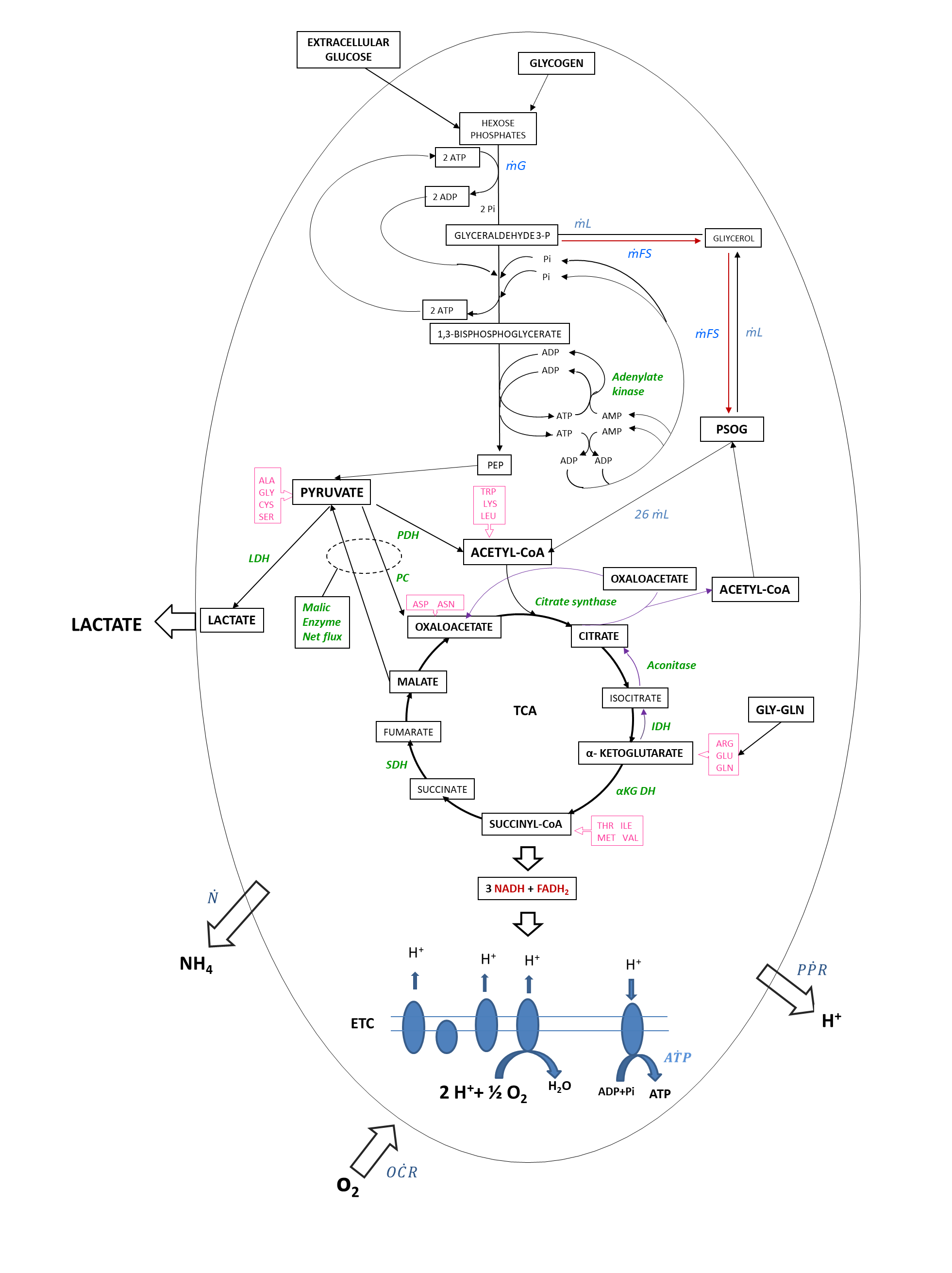

Catabolic pathways are represented with black arrows (glucose, amino acids and lipids. The reference PSOG (palmityl-stearyl-oleyl glycerol) triglyceride was the reference fat for lipid catabolism. The fluxes are represented in blue. Amino acids (in pink) are shown at the points of entry into the metabolic pathways. Enzymes are named in green. Purple arrows represent lipid synthesis pathway. (NADP= nicotinamide adenine dinucleotide phosphate, PPP= pentose phosphate pathway, PSOG= palmityl-stearyl-oleyl glycerol, Pi= inorganic phosphate, PEP= phosphoenolpyruvate carboxylase, PDH= pyruvate dehydrogenase, LDH= lactate dehydrogenase, PC= pyruvate carboxylase, IDH= isocitrate dehydrogenase, α-KGDH= α-ketoglutarate dehydrogenase, SDH= succinate dehydrogenase, GLN= glutamine, ASP= aspartate, ASN= asparagine, VAL= valine, ILE= isoleucine, ALA= alanine, ARG= arginine, CYS= cysteine, GLY= glycine, GLU= glutamate, MET= methionine, SER= serine THR= threonine, LEU= leucine, LYS= lysine, TRP= tryptophan, ṁG = irreversible glucose utilization flux, ṁFS = fat synthesis flux, O$\dot{C}$R= oxygen consumption flux, P$\dot{P}$R= proton production flux, $\dot{N}$= NH_4_ flux, A$\dot{T}$P= ATP flux, ETC= electron transport chain).
